# When precise expectations lose: Precision and Competition Among Hidden Causes in Voice Recognition

**DOI:** 10.64898/2026.06.10.731404

**Authors:** Carina Ufer, Fabian Schneider, Helen Blank

**Author notes:** Contributing authors.

## Abstract

Perception is often described as Bayesian inference, where precise expectations are assumed to dominate perception by pulling estimates strongly toward their mean. However, this effect may reverse when the hidden cause of the sensory input is unknown and must itself be inferred among competing alternatives. We tested this prediction in a voice-recognition task in which listeners learned three voices of speakers that differed in precision, defined as the inverse variance of the acoustic features of their voices. When classifying ambiguous utterances, listeners across five experiments systematically favored speakers with lower-precision, i.e., more variable voices, an effect that was strongest under high sensory ambiguity and amplified by explicit knowledge of speaker variance. Computational modeling showed that listeners tracked and used idiosyncratic, speaker-specific precision estimates that increased with learning accuracy and were stable across sessions. These findings show that precision does not simply determine the strength with which perceptual estimates are drawn toward a prediction; when the hidden cause must be inferred among competing alternatives, it also determines which prediction is favored. Our results identify precision as a critical determinant of perceptual inference when multiple hidden causes provide competing predictions about a stimulus.

## 1 Introduction

Perception is commonly conceptualized as hierarchical Bayesian inference, where the brain continuously estimates the most probable hidden causes of noisy sensory inputs by integrating sensory evidence with prior expectations [1–4]. In a hierarchical generative model, beliefs about hidden causes at higher levels generate probabilistic predictions about representations at lower levels. These top-down predictions constitute prior beliefs (i.e., expectations held prior to any sensory input) and are integrated with sensory evidence in a precision-weighted manner, such that the relative influence depends on their respective precision, i.e., inverse variance [5, 6]. Despite the central role attributed to precision in Bayesian and predictive-processing accounts, the role of prior precision remains comparatively poorly understood, partly due to inconsistent definitions and operationalizations [7]. Some accounts apply precision weighting only to prediction errors, without assigning precision to the prior itself [7–9]. Others describe “strong” or “reliable” priors as precise, while equating precision with prior probability, hence the probability assigned to a categorical hypothesis [10, 11] rather than the inverse variance of the predicted sensory states. Still others treat the presence of a prediction, relative to the lack thereof, as evidence of higher prior precision [12]. Direct manipulation of expected precision, i.e., the inverse variance of the expected features, has therefore been rare (but see [13, 14]).

Precision may affect inference differently depending on whether the hidden cause is known or must itself be inferred. When the cause is known (e.g., your phone shows the caller ID of your mother), inference concerns feature estimation: the brain uses prior expectations over her vocal features to interpret the incoming signal. Here, increasing prior precision relative to that of the sensory evidence gives the prior greater weight, thereby pulling perception more strongly toward the prior mean (Figure 1 A & B; [13, 14]). For example, when the prior over a speaker’s vocal features is highly precise, sensory input that deviates from the expected features barely shifts the perception away from that expectation.

**Fig. 1.**
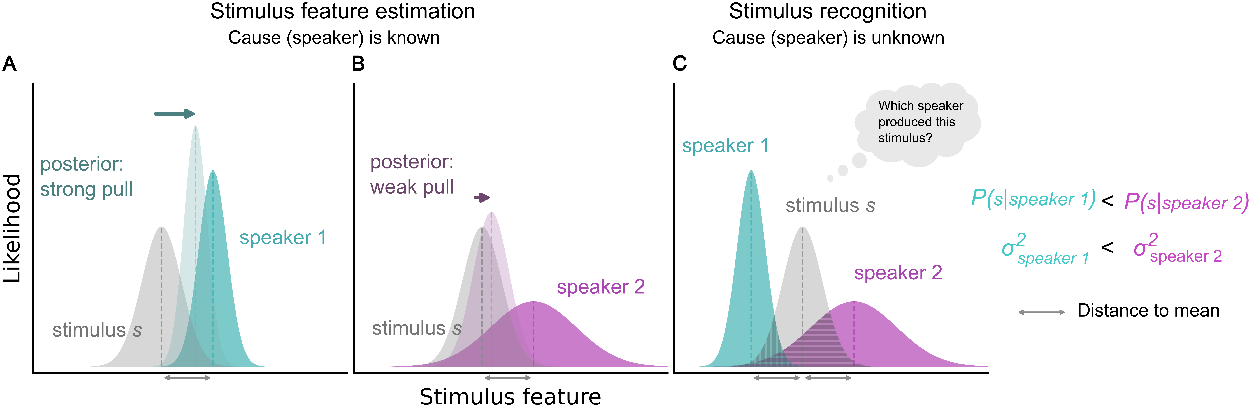
The effect of precision in feature estimation of cause inference. Speaker 1 has higher precision (i.e., lower variance) than speaker 2. When the category is known, high precision pulls the posterior more strongly towards the cause’s mean (**A**) than when precision is low (**B**). When the cause is unknown and the ambiguous stimulus *s* lies at an equal distance from the means of both distributions, the likelihood of *s* is lower under speaker 1 than under speaker 2, reflecting the reversal of the effect of precision in feature estimation or hidden cause inference.

When the hidden cause is unknown, however, inference involves an additional hierarchical level: cause inference, for example, determining who is speaking. For complex, real-world problems, such as identifying an unknown caller, the space of potential hidden causes (i.e., speakers) is effectively infinite. Each candidate cause is associated with a distribution of expected sensory features. These learned expectations are referred to as empirical priors [2]. When the cause is known, this distribution serves as the prior during feature estimation. When the cause is unknown, the same distribution functions as a conditional likelihood during cause inference. In this case, the system evaluates how probable the observed feature is under each candidate cause’s generative model. Crucially, the effect of precision reverses between feature estimation and cause inference. Increasing precision of a cause concentrates probability density near its mean, while reducing probability for stimuli farther away. Thus, although greater precision strengthens the influence of the predicted mean during feature estimation, it can reduce support for a candidate cause during cause inference when the observed features deviate sufficiently from predictions (Figure 1C). This apparent reversal follows from the different computational roles of precision in the two inference tasks. Although it follows directly from the mathematical structure of Bayesian inference, it has received little systematic experimental attention within predictive-processing frameworks.

Most experimental paradigms consider only a single cause and therefore do not directly test how precision shapes inference when several competing causes are possible. Although work on competing lexical hypotheses has examined inference over multiple candidate causes [15], the role of precision in such multi-cause settings remains unresolved.

Voice recognition provides an inherently hierarchical domain for investigating these computational principles. Identifying a speaker requires inferring the most likely hidden cause of an utterance. This inference is behaviorally relevant, as recognizing who is speaking shapes how speech is interpreted and how responses are selected [16–18]. Although listeners can typically recognize speakers effortlessly [19], individual utterances from the same speaker can vary substantially, such that no two instances are acoustically identical, making inference non-trivial [20–23]. Critically, speakers differ systematically in their degree of within-speaker variance, providing a principled basis for manipulating the precision of their speaker expectations [21, 24, 25]. We hypothesize that repeated exposure to a speaker’s voice gives rise to a learned empirical prior over that speaker’s acoustic features, which we refer to as the speaker representation. We propose that speaker representations incorporate not only the mean of these features [26] but also the precision. We further propose that voice recognition requires inferring which of several speakers most plausibly generated an utterance. To test this hypothesis, we manipulated the variance of speakers’ acoustic features and examined whether responses to ambiguous voice stimuli, positioned between learned speaker distributions, reflect Bayesian inference based on the empirical priors mean and variance (Fig 2A–C), or whether listeners’ decisions rely only on the empirical priors mean, disregarding precision (Fig 2D–F). Consistent with theoretical predictions, precision influenced inference: increasing the precision of a speaker’s voice reduced the probability that this speaker was inferred as the hidden cause of an ambiguous utterance. These findings demonstrate that precision plays opposite but complementary roles in feature and cause estimation, weakening inference when causes compete but sharpening perception when the cause is known.

**Fig. 2.**
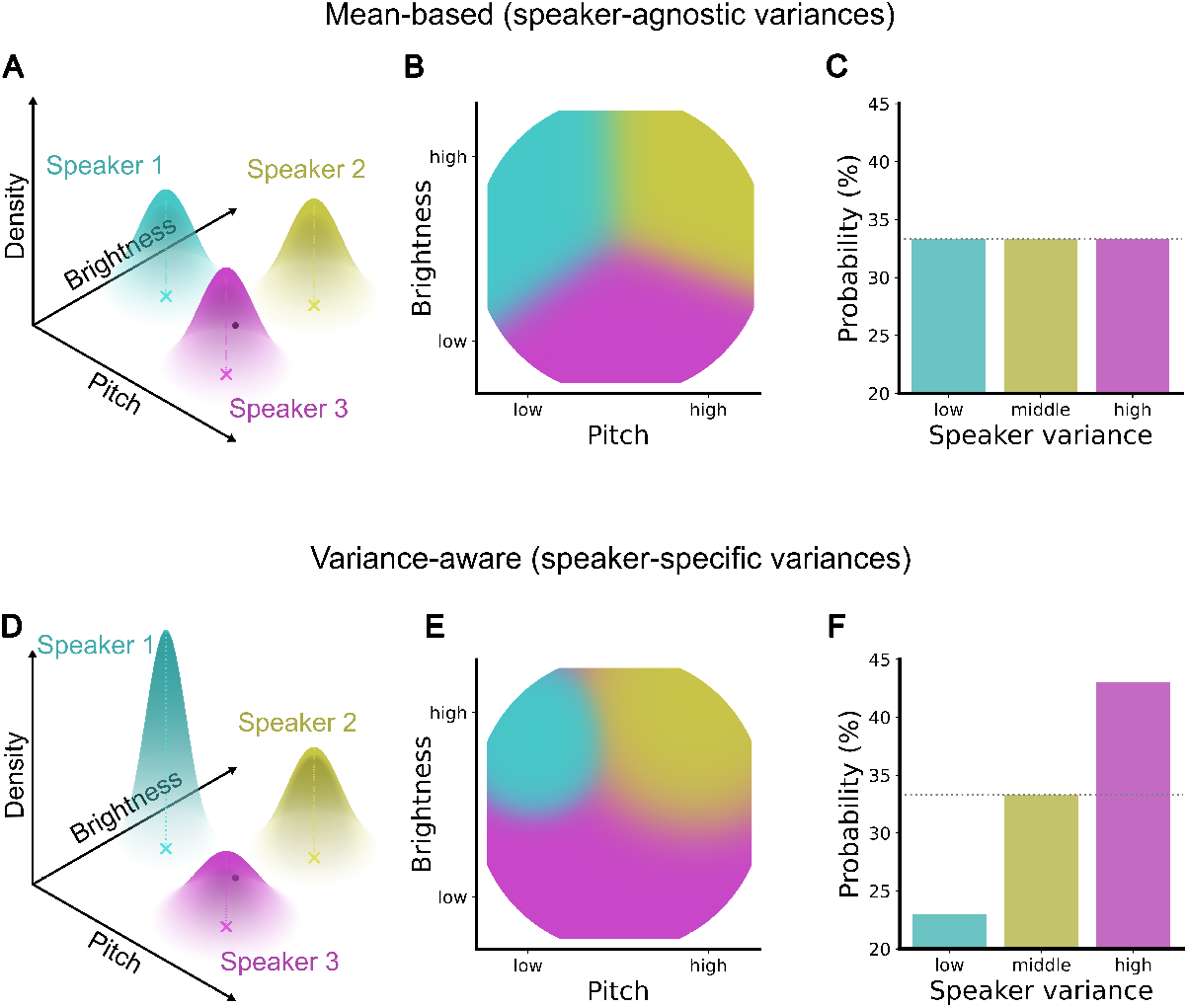
Hypothesis whether speaker variance is incorporated in the inference process or disregarded. **A** Three speaker representations with identical variance represented within a multidimensional voice space. **B**Predicted response pattern across the full feature space when inference over speakers is based on the means with equal variance; responses are determined by the stimulus’ distance to the speaker representations’ means. **C** Predicted average response pattern under identical-variance speaker representations, resulting in a uniform selection probability across voices. **D** Three speakers with distinct variances represented within the multidimensional voice space. **E** Predicted response pattern across the full feature space when speaker representations encode both the means and speaker-specific variances. **F** Predicted average response pattern assuming speaker-specific variances; voices with broader (high-variance) speaker representations exhibit a higher likelihood for ambiguous stimuli and are therefore expected to be selected more frequently, whereas voices with narrower (lower-variance) representations are expected to be selected less frequently.

## 2 Results

Across five online experiments, we tested whether perceptual inference over competing latent causes is influenced by the precision (operationalized as variance) of empirically learned priors. Participants learned the voices of three speakers (see Figure 3) defined by their mean and variance in a two-dimensional voice space spanned by two key acoustic features, voice pitch and brightness [25, 27, 28]. The three speakers had equidistant means but systematically manipulated variances (low, middle, high; see example participant in Figure 4A). The mapping between speaker identity and variance condition was counterbalanced across participants, such that each speaker identity was assigned to the low-, middle-, and high-variance conditions equally often. Each speaker representation corresponded to a distribution of voice exemplars spanning perceptually valid acoustic variation [20, 29–31], which participants learned to associate with the respective speaker identities. In contrast to two-alternative forced-choice paradigms, which can be solved by learning a single speaker and evaluating other stimuli relative to it, our design required participants to internalize full distributions of each speaker. This enabled direct investigation of how precision influences Bayesian inference over multiple competing hidden causes. Following learning, participants classified stimuli sampled from the full voice space, including regions of high uncertainty between speaker distributions. On each trial, participants inferred which speaker was the most likely hidden cause of the observed utterance. If inference incorporates precision in a Bayesian manner, ambiguous stimuli should be more frequently attributed to lower-precision (higher-variance) speaker representations due to their greater tolerance for deviation from the mean (see Figures 1 and 2E/F).

**Fig. 3.**
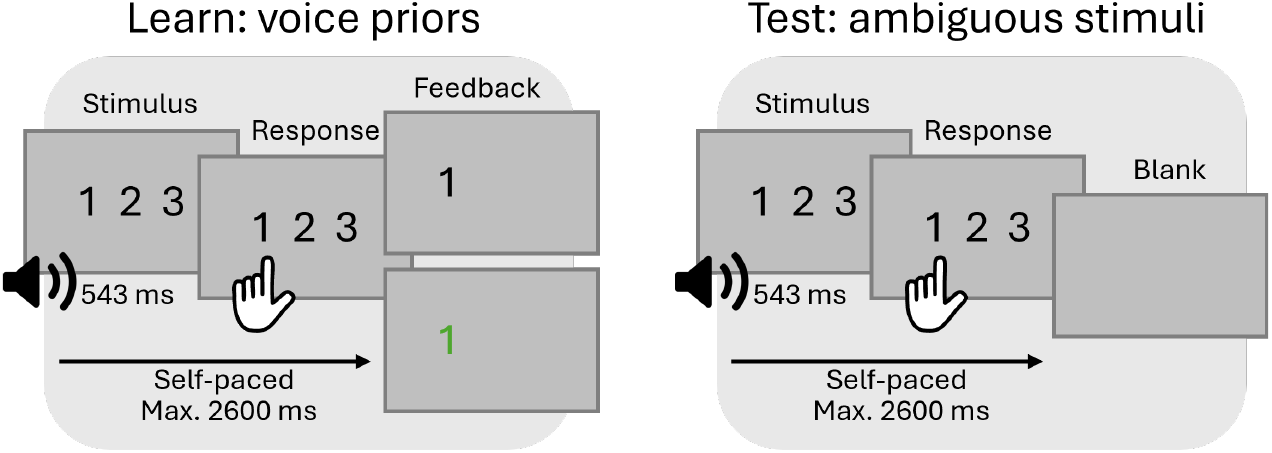
Schematic overview of the experimental procedure and analysis pipeline. Participants first learned the voices of three speakers, defined by their mean and variance in a two-dimensional voice space, by being presented with auditory stimuli and receiving feedback. On each learning trial, a spoken pseudoword (543 ms) was presented, and participants selected one of three speakers. Responses were self-paced within a maximum response window of 2600 ms and were followed by feedback indicating the correct speaker (500 ms). Learning and test blocks alternated across six blocks. Following learning, participants were presented with acoustically ambiguous test stimuli sampled from the same two-dimensional voice space. Participants responded using the same response options as in learning trials, but no feedback was provided; responses were followed by a blank interval (700 ms).

**Fig. 4.**
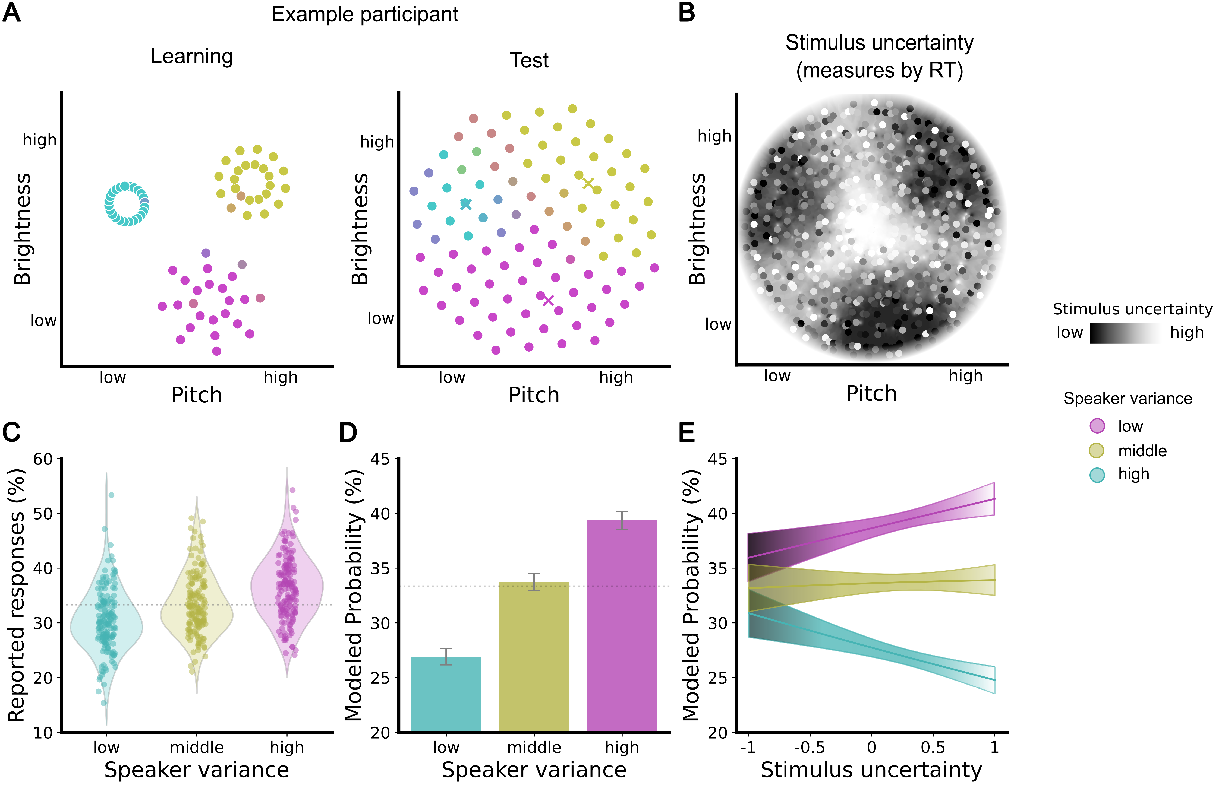
The influence of expected variance of a speaker’s voice on the inference of hidden causes under ambiguity. **A**. Example participant. Responses colored by proportion of responses for each speaker for the learning stimuli with feedback and the test stimuli without feedback. **B**. Stimulus uncertainty, measured by the average reaction time, shows that middle stimuli are more uncertain than stimuli closer to the centers of the speakers. **C**. Proportion of reported responses per speaker (raw data). Single dots represent participants. **D**. Probability of assigning each speaker as the hidden cause of an utterance based on the coefficients of the multinomial logistic model. The bars depict the effect independent of stimulus features over all experiments. Error bars represent the 95% confidence intervals. **E**. Probability of assigning each speaker as the hidden cause of an utterance based on the coefficients of the multinomial logistic model. The influence of speaker variance increases with stimulus uncertainty: the greater the uncertainty about the stimulus, the more likely it is to be attributed to the high-variance speaker.

Data were pooled across five online experiments (N = 260; see Table 1 for specific experimental variations), sharing a similar learning–test structure. To ensure that participants adequately learned the speaker representations to apply them to ambiguous test stimuli, we preregistered an inclusion criterion of *≥* 80% accuracy during learning (similar to [32]), yielding 162 above-criterion participants. All effects were corroborated in the full sample (Supplementary Tables S1 – S8).

**Table 1.** Experimental designs over experiments.

| | Variance instruction | None-option as response | Test stimuli | Repetition of experiment | N | $n_{\text{above-criterion}}$ |
| --- | --- | --- | --- | --- | --- | --- |
| Experiment 1 | explicit | no | separate block | no | 30 | 18 |
| Experiment 2 | explicit | yes | separate block | no | 27 | 18 |
| Experiment 3 | explicit | no | intermixed | no | 34 | 18 |
| Experiment 4 | explicit | no | intermixed | yes | 21 | 14 |
| Experiment 5 | implicit | no | separate block | no | 169 | 108 |
| Total |  |  |  |  | 260 | 162 |

### 2.1 Expected variance of a speaker’s voice influences the inference of the hidden cause

To test whether hidden cause inference is modulated by precision in a multi-cause setting, we fit a multinomial model predicting participants’ responses. Because voice-to-variance mappings were counterbalanced across participants, responses were standardized to a common latent precision space, ensuring inference was evaluated independently of specific speaker identities. The model thus captured selection probabilities over competing latent causes as a function of the expected precision of each speaker representation (response: low, middle, high variance speaker).

The results revealed a robust bias in hidden cause inference driven by precision. Participants were most likely to attribute ambiguous utterances to speakers with lower precision representations (higher variance, Figure 4D). Independent of stimulus features, participants attributed utterances to the high-variance speaker most often (39.4%, CI [38.6–40.2%]) compared to the middle (33.7%, CI [32.9–34.5%]) and low-variance speakers (26.9%, CI [26.2– 27.7%], all *p <* .001; Table S9 and S10). Thus, under sensory uncertainty, inference over competing latent causes is systematically biased toward internal representations with lower precision.

### 2.2 The effect of expected variance is strongest when sensory input is uncertain

We next tested whether the influence of speaker precision on hidden cause inference depends on sensory uncertainty. Observers should be more certain about the latent cause of a stimulus the more similar the features of the stimulus are to those of the mean of a speaker’s voice. According to Bayesian accounts, the influence of precision should be strongest for highly ambiguous sensory inputs. Mean reaction time (RT), averaged across participants per stimulus, was used as a fine-grained measure of stimulus ambiguity, rather than distance from the means of the speaker representations. This is because features that are farther from the center of the vocal space along the direction of a speaker’s mean are perceptually exaggerated, leading to caricature-like samples that are easier to classify (see Methods) [33]. The multinomial model confirmed that the effect of precision varied systematically with stimulus uncertainty (Figure 4E, Tables S11 and S12). When stimulus uncertainty was lowest, no bias toward the lower-precision speaker was observable, as all confidence intervals approached 33% (low-variance speaker: 30.9% CI [28.7–33.1%]; middle-variance speaker: 33.2% CI [31.0–35.3%]; high-variance speaker: 35.9% CI [33.8–38.1%]), suggesting that participants could correctly infer the intended speaker. In contrast, when stimulus uncertainty was high, participants were more likely to infer the high-variance speaker as the hidden cause (41.3% CI [39.8–42.8%]) compared to the middle-(33.9% CI [32.5–35.3%]) or low-variance speaker (24.8% CI [23.6–26.0%]).

### 2.3 The effect of expected variance is stronger with explicit knowledge but persists with implicit learning

Next, we investigated whether the influence of precision on hidden cause inference depends on the degree of explicit knowledge and learning of the speaker representations. Participants either learned the speakers implicitly (Experiment 5) or received explicit information about the variance of each speaker’s distribution while learning the speakers (Experiments 1–4). The effect of precision on the inference of hidden causes was present under both implicit and explicit learning, but was significantly amplified by explicit knowledge (Figure 5A). Under implicit learning, participants most frequently attributed stimuli to the high-variance speaker (37.8%, CI [36.9–38.7%]), followed by the middle-variance (34.0%, CI [33.1–34.9%]) and low-variance speakers (28.1%, CI [27.3–29.0%]). Explicit knowledge further increased attribution to the high-variance speaker (+4.0%) and reduced attribution to the low-variance speaker (-3.1%; Tables S13 and S14), indicating an amplification of the precision-dependent inference effect.

**Fig. 5.**
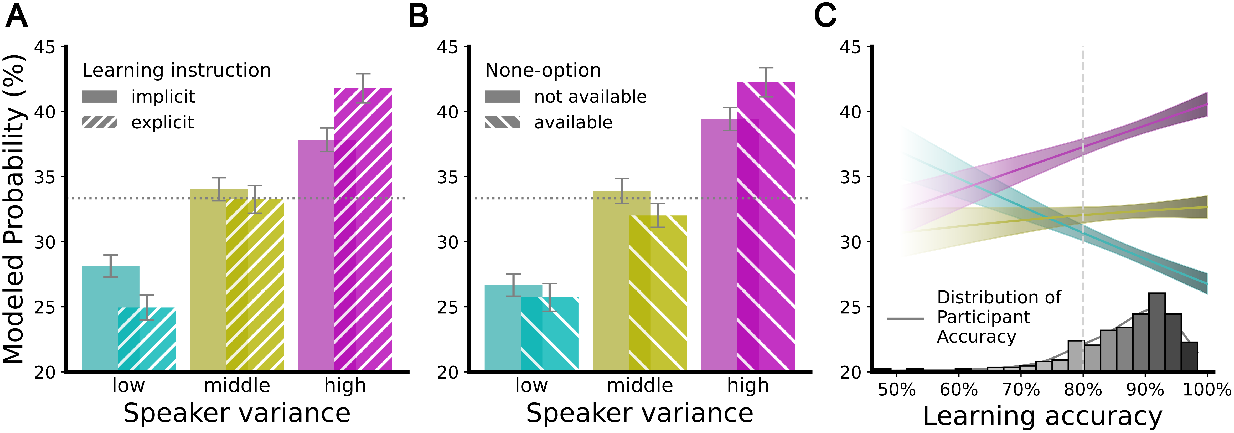
The influence of experimental conditions and learning accuracy on the inference of a speaker as the hidden cause. **A**. The effect of variance on the inference of the hidden cause is amplified with explicit (Experiments 1–4) knowledge about the variances but persists when variances are implicit (Experiment 5). **B**. The effect of variance on the inference of the hidden cause persists when participants were given the option to respond that “None of the speakers” is the hidden cause (Experiment 2). **C**. The influence of variance is amplified by higher learning accuracy in the learning blocks.

To test whether the observed amplification of the effect by explicitness was genuinely due to a Bayesian inference process rather than an explicit response bias, we investigated whether the effect persisted when participants were given the option to respond that none of the speakers was the hidden cause of an utterance (Experiment 2). However, the none-option was rarely used (2.1% *±* 3.4%), and its availability did not reduce the effect of precision (Figure 5B, Tables S15 and S16). The bias toward the high-variance speaker remained robust (42.3%, CI [40.1–44.4%]) relative to the middle-(32.0%, CI [30.0–34.0%]) and the low-variance speaker (25.7%, CI [23.9–27.6%]).

### 2.4 The effect of expected variance scales with learning accuracy

We reasoned that the influence of precision on hidden-cause inference should depend on how accurately participants learned the intended experimental manipulation of speaker-specific precision. We tested whether the effect of precision scaled with learning accuracy by including all participants regardless of their learning performance. Consistent with our hypothesis, the effect of precision on hidden-cause inference amplified with higher learning accuracy (Figure 5C, Tables S17 and S18).

Participants with lower learning accuracy (*−*1 SD) showed a weaker bias toward the high-variance speaker (36.9%, CI [36.2–37.6%]) compared to those with higher learning accuracy (+1 SD; 39.8%, CI [39.1–40.5%]). Similarly, participants with lower learning accuracy showed a reduced bias towards the low-variance speaker (31.1%, CI [30.5–31.8%]) compared to participants with high learning accuracy (27.7%, CI [27.0–28.3%]). Thus, the influence of precision on hidden-cause inference scaled with learning accuracy, such that more accurate learning was associated with a stronger precision-dependent selection of the inferred cause. With greater learning accuracy, the tendency to infer the low-variance speaker as the hidden cause decreased, while the tendency to infer the high-variance speaker as the hidden cause increased.

## 3 Computational model to estimate idiosyncratic precision in speaker representations

The multinomial model showed that the effect of precision-dependent inference was stronger in participants with higher learning accuracy. This raises the possibility that participants with lower learning accuracy acquired idiosyncratic speaker representations that deviated from the experimentally intended distributions, while still performing Bayesian inference relative to these internal representations.

Since the multinomial model cannot provide insights into the underlying cognitive mechanism driving hidden-cause inference, we developed and compared computational models capturing different alternative cognitive mechanisms underlying hidden-cause inference. Our first objective was to determine whether participants incorporate model-specific precision by testing whether models including speaker-specific variance better explain behavior. Our second objective was to assess whether allowing interindividual differences improves model fit, thereby revealing idiosyncratic, participant-specific speaker representations.

We therefore compared four candidate models of different underlying cognitive mechanisms varying along two dimensions: whether model-specific precision was incorporated (speaker-agnostic variance vs. speaker-specific variance) and whether speaker representations were shared (general) or participant-specific.

Across models, Bayesian inference was formalized as the likelihood of each test stimulus under each speaker’s distribution, determined by its distance to the mean of the empirical speaker prior weighted by the variance. Smaller variance parameters produced sharper category boundaries, whereas larger variance parameters produced broader, imprecise category boundaries.

### 3.1 Individuals incorporate idiosyncratic variance in speaker representations

We first confirmed the speaker-specific variance model under conditions in which participants had reliably learned the intended speaker distributions. In the above-criterion sample, the speaker-specific variance model outperformed the speaker-agnostic variance model, and the estimated variance parameters recovered the experimentally imposed ordering (low to high variance; Figure 6B, Tables S19 and S20). Thus, when speaker representations are well learned, their precision is incorporated into hidden-cause inference.

**Fig. 6.**
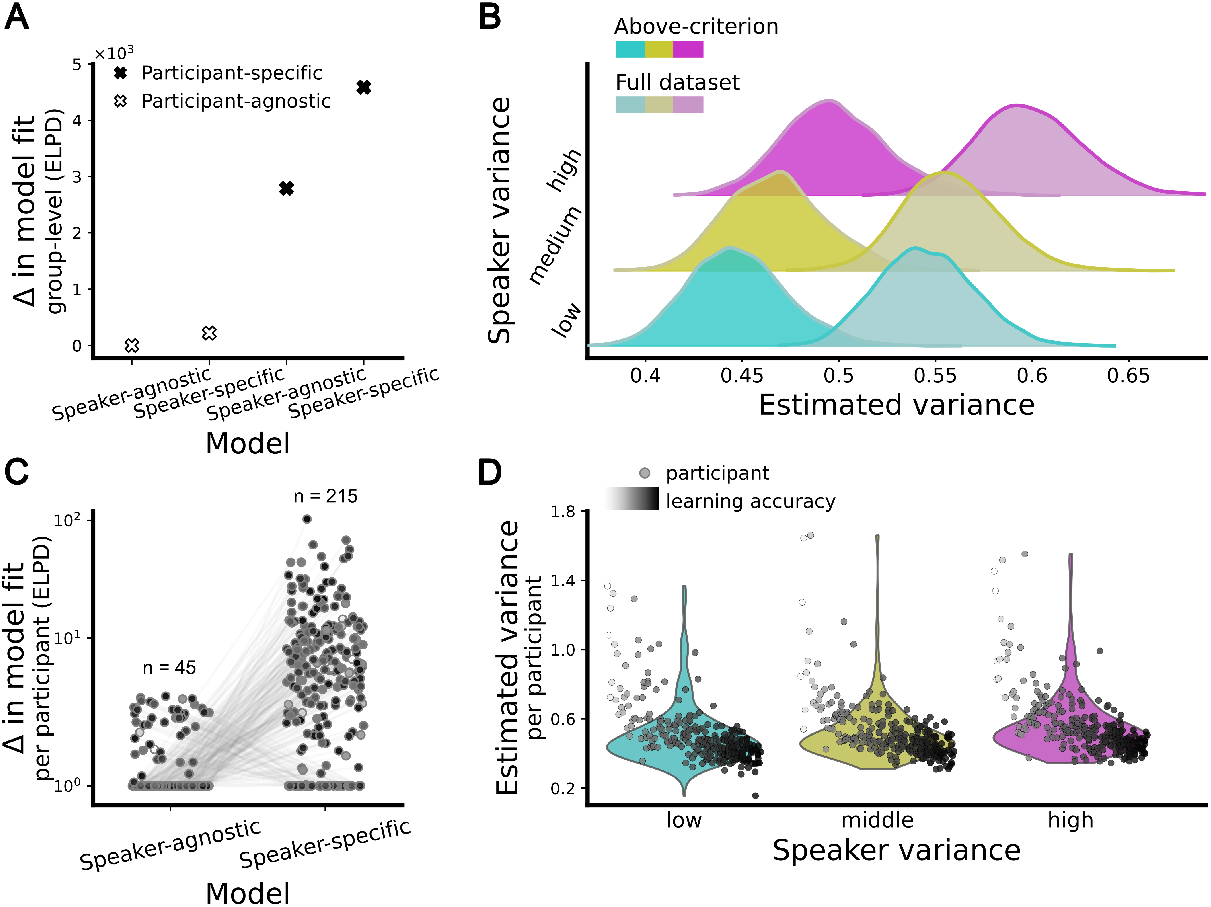
Computational model results and voice variance estimates. **A**. Differences in the expected log-pointwise predictive density (ELPD) relative to the value of the baseline model, illustrating model performance across the full dataset. Higher ELPD values indicate better predictive accuracy. The model incorporating distinct variance estimates for each voice together with a participant-specific estimate is superior to the other models. **B**. Posterior distributions of the group-level speaker-specific variance parameters from the computational hierarchical model, shown separately in different color shades for the above-criterion and full datasets. The estimated parameters are ordered according to the expected variance. **C**. Individual-level differences in expected log-pointwise predictive density between the speaker-agnostic variance and speaker-specific variance model, computed for each participant separately. Most participants’ responses (215 of 260) are best explained by the variance model, indicating its superior predictive performance at the individual level. **D**. Estimated variance parameters for each participant derived from the hierarchical variance model. Each dot represents an individual participant’s variance estimate. The grey shading indicates the participant’s learning accuracy from the separate learning blocks, with lower accuracy associated with higher variance estimates, suggesting less precise boundary delineation between the voices.

Participants with lower learning accuracy, however, may have acquired idiosyncratic speaker representations that deviate from the intended distributions, while still performing Bayesian inference relative to these internal representations. To test this, we repeated the analysis in the full dataset, including all participants irrespective of learning performance.

In this more heterogeneous sample, the speaker-specific variance model again outperformed the speaker-agnostic model (Figure 6A, Table S21), with a substantial improvement in predictive accuracy (Δ expected log pointwise predictive density (ELPD) *>* 200; differences *<* 4 are considered small; [34]), indicating that precision continued to shape inference.

We next tested whether allowing idiosyncratic, speaker-specific precision improves model fit. Introducing participant-specific speaker-specific variance estimates to the models resulted in a substantial improvement in model fit (Δ ELPD *>* 2000). In line with our hypothesis, the participant-specific speaker-specific variance model provided the best overall fit. Model comparison at the individual participant level further confirmed the superiority of the speaker-specific variance model, which outperformed the speaker-agnostic variance model (215/260 participants). This provides evidence that participants used speaker representations with idiosyncratic variances during inference.

Group-level parameter estimates in the full dataset still followed the expected ordering, though less distinctly (Figure 6B, Table S22). The estimated *σ* was lowest for the low-variance speaker (0.55, 95% highest density interval (HDI) [0.50, 0.59]), followed by the middle-variance speaker (0.56, 95% HDI [0.51, 0.61]) and the high-variance speaker (0.60, 95% HDI [0.55, 0.65]). This suggests that participants broadly learned and applied the intended variance structure, while exhibiting systematic individual deviations. This is further supported by the stronger group-level effect observed in the above-criterion sample compared to the full dataset, indicating that participants who failed to reach the learning criterion acquired speaker-specific variances that deviated from the experimentally intended structure.

To further characterize these deviations, we grouped participants according to the ordering of their idiosyncratically estimated speaker-specific variances derived from the computational model (Figure 7A). Especially participants who fell below the preregistered accuracy criterion demonstrated deviations from our intended variance pattern (Figure 7B), indicating that they were more likely to acquire alternative variance structures.

**Fig. 7.**
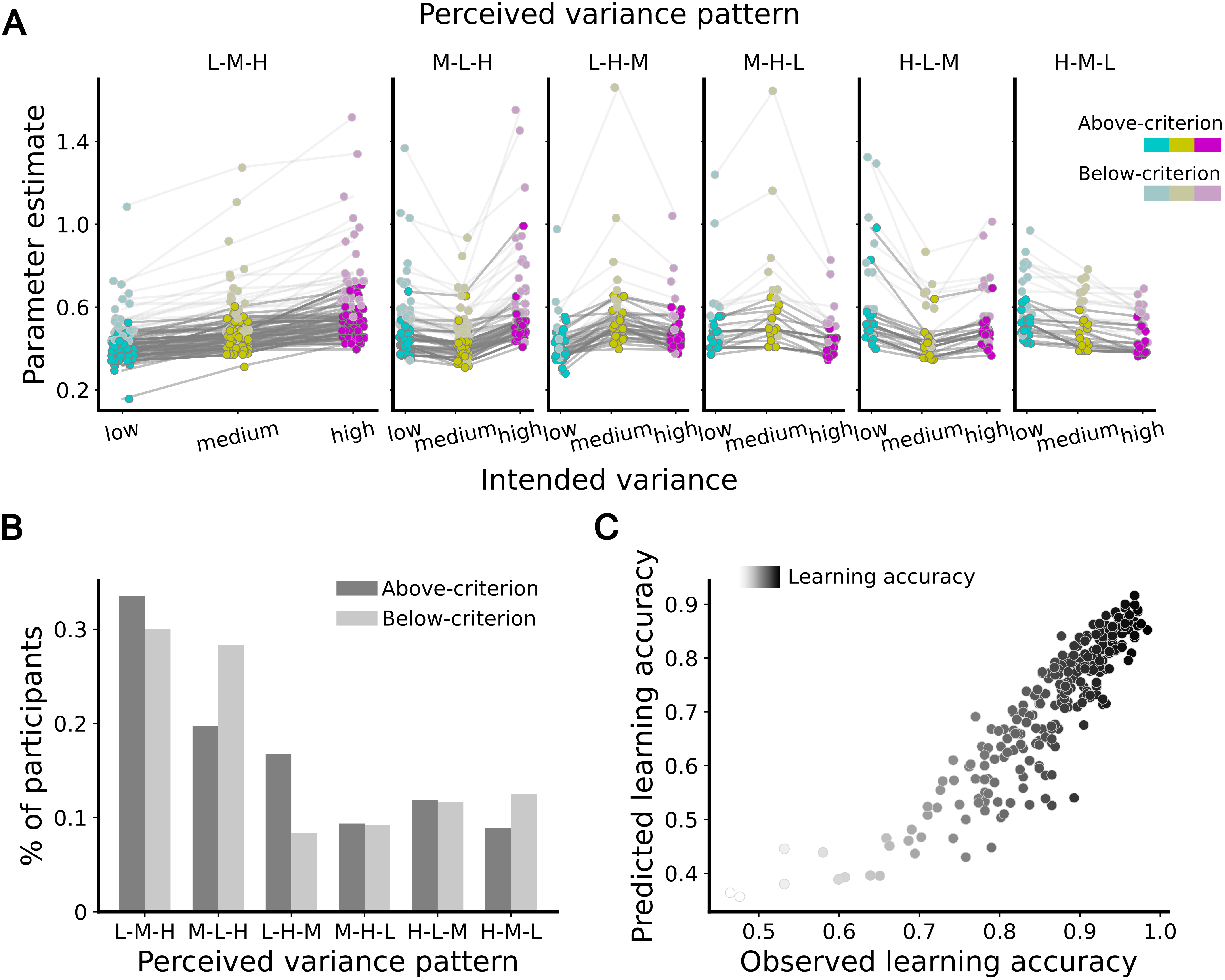
Interindividual differences in variance of speaker representations. **A**. Participants are grouped according to the order of their idiosyncratic variance estimates. Group 1 represents the expected ranking: the low-variance speaker has the lowest variance estimate, followed by the middle-variance speaker, and the high-variance speaker with the highest estimate. Bright colors indicate above-criterion participants, while lighter colors denote below-criterion participants. **B**. Counts of participants within each variance group, separated by validity status (above-criterion vs. below-criterion). **C**. Correlation between the learning accuracy predicted by the winning hierarchical variance model and the observed learning accuracy across participants.

Together, these results show that perceptual inference over competing latent causes operates over idiosyncratic, hierarchically structured internal representations which incorporate precision.

### 3.2 Idiosyncratic variance in speaker representations is explained by manipulated variance, learning accuracy, and perceived variability

Having established that participants rely on their idiosyncratic speaker-specific variances that may systematically deviate from the experimentally intended structure, we next investigated which factors explain these individual differences.

As a first, confirmatory step, we tested whether the experimentally intended variance of the speakers could predict participants’ idiosyncratic variance estimates. Consistent with the group-level ordering of variance estimates (Figure 6B), a linear mixed-effects model showed that idiosyncratically acquired variance in the speaker representations systematically increased with the intended variance (*β* = 0.027, *p < .*001; Table S23). This confirms that participants, on average, encoded the experimentally defined variance structure in their internal representations. Having established this correspondence, we next investigated whether learning accuracy explains deviations from the intended variance structure. At the group level, variance estimates obtained from the full sample were markedly higher than those derived from the above-criterion sample alone (Figure 6B). Because the only difference between these datasets is the inclusion of below-criterion participants, this pattern suggests that lower-performing participants contributed to the increase in estimated variance. Including standardized mean learning accuracy as a predictor of the idiosyncratic variance estimates revealed a negative relation (*β* = *−*0.131, *p < .*001), indicating that participants with lower learning accuracy acquired speaker representations with higher variance. In computational terms, higher variance reflects broader and less well-defined category boundaries for a voice. Thus, lower learning accuracy is associated with less precise internal representations of speakers, which are then carried forward into the inference stage.

We further examined whether additional individual differences explain idiosyncratic variances in speaker representations. Incorporating participant characteristics and their subjective perceptions of voice features into the linear mixed-effects model revealed that subjectively perceived variability in a voice significantly predicted idiosyncratic variance estimates (*β* = 0.013, *p < .*01; Table S24). This suggests that speaker representations are shaped by subjective perceptual experiences and interpretations rather than solely by the true acoustic features. Listener age showed a small positive effect (*β* = 0.003, *p < .*01), however, the magnitude of this effect seems negligible. None of the other perceived voice characteristics (familiarity, distinctiveness, and naturalness), nor listener sex, significantly predicted idiosyncratic variance estimates (all *p > .*05, Table S24). Overall, these findings indicate that individual perceptions during learning play a crucial role in the formation of empirical priors.

### 3.3 Idiosyncratic speaker representations predict learning accuracy

We next tested whether participants’ idiosyncratic variance estimates reflect how they perceived and learned the stimulus distributions. If speaker representations are shaped by subjective perception rather than true stimulus statistics, then systematic misperception during learning should give rise to empirical prior distributions that deviate from the intended distributions but still govern subsequent inference.

Ideally, the parameters of the speaker representation would be estimated directly from learning behavior and used to predict responses at test. Since the learning stimuli were designed to reflect the intended variances, were limited in number, and did not include highly ambiguous samples, which prevented a reliable estimation of variance parameters from the learning phase alone, we employed a reverse modelling approach to reconstruct each participant’s individual learning accuracy from their idiosyncratic variance estimates obtained from the computational model. Specifically, we predicted responses to the learning stimuli by sampling from the posterior distribution of the model and derived predicted learning accuracy. If the model-estimated variances capture the internal representations acquired during learning, they should reproduce participants’ actual learning accuracy.

We found that model-predicted accuracy per participant closely matched the observed performance (*r* = 0.90, *p < .*001; Figure 7C). This strong correspondence indicates that participants’ behavior is well explained by the idiosyncratic speaker representations they acquired, supporting the interpretation that perceptual inference operates over internally learned idiosyncratic precision, even when these deviate from the experimentally intended distributions.

### 3.4 Idiosyncratic speaker representations are stable within a participant

As subjective perception during learning influenced how speaker representations were formed, we hypothesized that these perceptual differences may be shaped by prior experiences (hyperpriors), potentially linked to interindividual differences in acoustic processing and voice recognition ability [35–38]. If hidden-cause inference is shaped by participant-specific hyperpriors, then these should be stable and should not change substantially after a single exposure to our stimuli. To test whether the idiosyncratic variance estimates reflected such stable hyperpriors, we invited participants for a second session. If participants brought their individual hyperpriors into the experimental setting, their idiosyncratic speaker representations should stay robust and their responses in the second session should be more accurately predicted by their own parameter estimates from the first session than by those of other participants.

For each participant, we compared second-session responses to predictions generated from all participants’ first-session idiosyncratic variance estimates. This yielded prediction scores quantifying how well each participant’s full set of first-session variance estimates accounted for their second-session behavior relative to others. Participants were then ranked according to how well their own idiosyncratic variance estimates predicted their responses compared to the estimates of other participants. Predictions based on participants’ own idiosyncratic speaker representations were significantly better than chance (one-sided Wilcoxon signed-rank test, *p < .*001; Figure 8A), indicating that behavior in the second session was best explained by participants’ own inferred internal speaker representations. To further quantify this effect, we used a hierarchical Bayesian model to estimate likelihood differences between self- and other-based predictions (self *−* other). Positive values indicate that responses are more likely under a participant’s own idiosyncratic speaker representations.

**Fig. 8.**
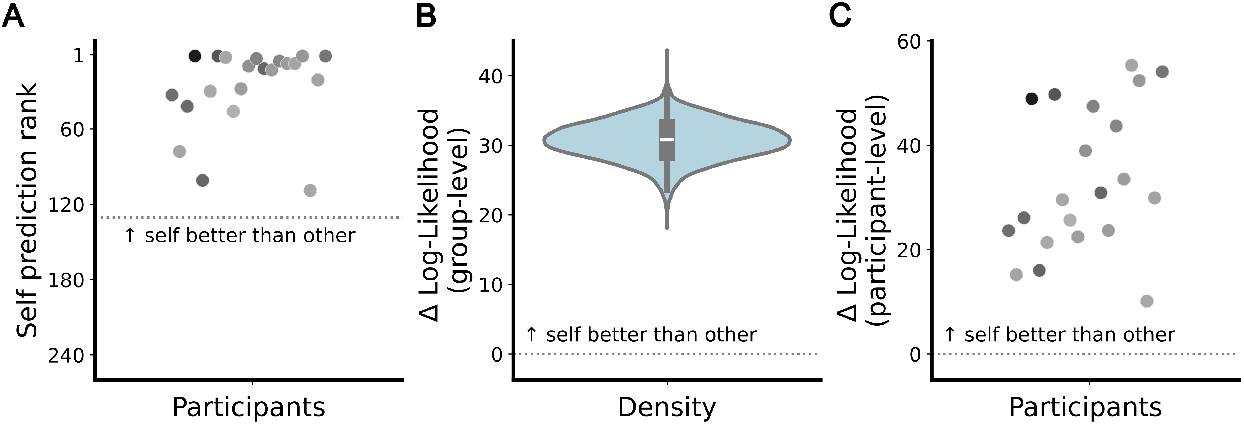
Stability of idiosyncratic structure in internal speaker representations. **A**. Rank of each participant’s prediction of their responses based on their own model parameters learned in the first run of the experiment. The ranks are significantly lower than the chance rank expected if there were no advantage in predicting responses using one’s own parameters relative to those of other participants. **B**. Group-level estimate of the self-prediction advantage, defined as the difference in log-likelihood between predictions generated from a participant’s own parameters and predictions generated from parameters of other participants. **C**. Participant-specific estimates of the self-prediction advantage, quantified as the difference in log-likelihood reflecting improved prediction of one’s own responses compared to predictions based on other participants’ parameters.

At the group level, this self-prediction advantage was positive (Figure 8B) and was consistently observed across all participants (Figure 8C). Together, these findings suggest that idiosyncratic hyperpriors shape participants’ subjective perception, which in turn leads to stable idiosyncratic variance structure in speaker representations.

## 4 Discussion

Our study provides empirical evidence that humans incorporate not only the mean but also the precision of learned acoustic feature distributions in their speaker representations, consistent with hierarchical Bayesian inference. Critically, our findings demonstrate the predicted reversal in precision’s role depending on whether the hidden cause is known or must itself be inferred. When a single candidate cause is assumed, higher precision is expected to pull the perceptual estimate more strongly toward the prior mean [13, 14]. We show, however, that once multiple hidden causes compete as candidate causes, the same precision parameter instead reduces a candidate’s posterior probability. A high-precision distribution assigns low likelihood to sensory input that deviates from its mean, whereas a low-precision distribution remains more tolerant. Consistent with this, participants preferentially attributed ambiguous stimuli to speakers with lower-precision (higher-variance), particularly under increased sensory uncertainty. These results provide empirical support for the dual but complementary role of precision in hierarchical generative models: the parameter that sharpens perception when the cause is known simultaneously reduces support for a candidate cause during cause inference when the cause is unknown.

Our results align with classic categorization research showing that ambiguous stimuli are preferentially assigned to more variable categories, e.g., an object of intermediate size is more likely to be categorized as a pizza (highly variable in size) than as a coin (narrowly constrained) [39]. Compared to previous studies that found the effect inconsistently [40, 41], we demonstrate a robust advantage for low-precision candidate causes. This may reflect that our study constitutes a genuine, ecologically valid cause attribution problem rather than an arbitrary similarity judgment over experimenter-defined categories, and that three competing speakers in a multidimensional acoustic space increased the inferential pressure to learn and distinguish their respective distributions.

Previous work proposed that greater precision, induced by learning, pushes estimates away from category boundaries via an implicit category decision, which they inferred from model fits rather than testing directly [42]. Our study used explicit categorization, and consistent with this account, as precision increased with learning, deviant stimuli became less likely under the precise speaker representations and were increasingly attributed to alternative speakers. Our results thus provide converging support for a mechanism in which learning-induced precision shapes categorical inference.

Previous work has shown that observers can learn both mean and precision of a single cause’s distribution and use these parameters during feature estimation [13, 14]. We extend this framework by showing that individuals can learn empirical priors for multiple competing hidden causes with distinct means and precisions, and use these parameters during cause inference.

By incorporating multiple competing hidden causes, our experimental design captured an ecologically relevant aspect that laboratory paradigms often overlook. In natural environments, sensory input is rarely evaluated against a single hypothesis. Instead, the perceptual system must infer which of several candidate causes generated the input, with each associated with its own probability distribution over sensory states. The idea that internal generative models comprise multiple competing predictions, as we show here for voice recognition, has also been demonstrated in language processing, where listeners must infer which of several possible words was spoken [15]. Although stimulus features typically vary continuously, such as colour or size, it has been proposed that these dimensions are discretized into categorical representations [43, 44], which can be interpreted as distinct candidate causes within generative models. Consistent with this view, recent neural evidence indicates that multiple candidate predictions are represented concurrently, with decoding accuracy improving as additional plausible hidden causes are incorporated as predictors [17]. Moreover, neural population codes represent information about the number of competing hypotheses, further corroborating the role of competing predictions in perceptual inference [45]. Our study not only implemented multiple hidden causes but also manipulated the precision over the expected sensory features, thereby advancing our understanding of how inferences about hidden causes are made in the face of multiple possible options with different distributions.

The presence of the effect under implicit learning indicates that precision is automatically extracted during the acquisition of empirical priors, consistent with previous findings showing that prior variance can be acquired through experience alone [13]. Providing explicit information about precision further amplified the effect, similar to previous work [40]. This suggests that while precision is encoded during perceptual learning, its influence can be strengthened with explicit information. In contrast, previous work showed that implicitly acquired prior probabilities exerted a stronger influence on perception than explicitly acquired prior probabilities [46]. In our study, the presence of the effect under implicit conditions rules out a post-perceptual decision bias, pointing to a genuine perceptual component with possible additional decisional amplification driven by explicit knowledge. Moreover, the effect persisted when participants could reject all candidate causes, indicating that it is not merely a consequence of a forced-choice strategy but reflects genuine perceptual inference.

Our results further show that perceptual inference operates over idiosyncratic speaker distributions that vary systematically across individuals. Hierarchical Bayesian modeling showed that participants internalized speaker-specific variances that partially deviated from the intended stimulus statistics, indicating individualized generative models. Together with previous work [17, 47, 48], this underscores the importance of participant-level modeling in perceptual inference. These idiosyncratic speaker representations were behaviorally relevant. Model-derived estimates predicted participants’ responses during learning, demonstrating that the speaker representations aligned with how sensory information was encoded. Moreover, participants with more accurate learning exhibited inference patterns more closely aligned with computationally predicted Bayesian inference patterns based on the intended precision structure.

We observed substantial interindividual differences in participants’ idiosyncratic priors. Beyond the experimental manipulation and learning accuracy, subjective perceptions of variability within a voice explained idiosyncratic estimates, illustrating that perception fundamentally involves a subjective interpretation of stimulus features. One explanation for these perceptual differences is that perception is shaped by hyperpriors formed through lifelong experience, influencing how acoustic stimuli are encoded. Such hyperpriors may underlie interindividual differences in processing voice-relevant acoustic features [36, 38] and in voice recognition ability [19, 49], giving rise to stable differences in the acquisition of speaker representations. Those hyperpriors are unlikely to and arguably should not change through exposure to our experiment. Consistent with this, idiosyncratic representations remained stable across sessions. Alternatively, idiosyncratic representations might arise from differences in how information is weighted, integrated, or updated during learning of empirical priors. Interindividual differences in statistical learning and in the use of prior information are well documented across perceptual domains [50–53], and can lead to systematic differences in prior acquisition.

Together, these findings suggest that interindividual differences in representations reflect trait-like differences in perceptual encoding shaped by hyperpriors, differences in predictive computations, or an interaction of both.

Our findings have important implications for voice perception. While previous studies show that representations of a speaker’s voice are shaped by statistical regularities and encoded as averages in a multidimensional space [26, 28, 33, 54], we demonstrate that speaker identity recognition depends not only on the mean but also on the precision of the internal speaker representation. Speaker-specific precision, reflecting the variance across utterances [20, 21, 23], provides a critical cue for hidden-cause inference under uncertainty and may support stable speaker identification despite natural fluctuations in vocal features. Consistent with this, listeners can reliably recognize speakers across variable utterances [22, 23, 26, 55], indicating that the perceptual system extracts information from sensory input to form expectations. Together, these findings suggest that speaker representations are formed through the integration of voice samples, capturing both the mean and variance.

In our study, all voices were deliberately equidistant from the population mean to insulate the influence of speaker precision. Future work could test whether the encoding of precision depends on proximity to the population mean. Previous work suggests that voice perception is structured relative to a population-level prior, with voices encoded as deviations from this reference [54]. Differences near the population mean are perceived more distinctly than equivalent differences in the tails [56], indicating non-uniform discriminability across the space. This perceptual warping raises the possibility that speaker-specific variance is also encoded non-uniformly across the space. A similar effect may arise within a voice: familiarity sharpens voice representations, with familiar voices perceived as more distinct than unfamiliar ones [57], hinting at possible experience-dependent modulation of perceived variance. In the present study, familiarity did not explain idiosyncratic precision estimates, possibly because familiarity ratings were generally low [58].

Our paradigm concerned the hidden-cause inference (inferring the speaker) and did not focus on feature estimation (inferring pitch and brightness). In natural listening, however, these processes are intertwined. Recognizing a speaker requires processing of vocal features, yet those features may in turn be perceived differently once the hidden cause is inferred, reflecting the differing roles of precision described in the introduction. Our experiment cannot determine how the brain resolves this interdependence. One possibility is a coarse-to-fine process in which initial processing of vocal features generates a first hypothesis about the speaker, which then shapes the interpretation of subsequent features by pulling them toward what that speaker typically sounds like, with both estimates iteratively refined across processing levels. Alternatively, the brain may never commit to a single speaker identity, but instead computes feature estimates as a weighted average over candidate speaker representations, with each speaker’s contribution weighted by its posterior probability given the current sensory input. The exact mechanism, and the processing stage(s) at which feature and cause inference interact, remains to be determined. Future work jointly manipulating uncertainty at both levels, combined with neural measures sensitive to vocal feature representations, will be needed to resolve this.

Although our experiment moves closer to naturalistic settings by including multiple competing hidden causes, it remains a tightly controlled paradigm. We defined each speaker by manipulating only two acoustic dimensions, pitch and brightness, based on a single artificial voice sample. Natural voices vary across additional features such as harmonicity, jitter, speech rate, intonation, and loudness [25, 59], which may all contribute to the formation of a speaker’s voice representations and hence to perceptual inference. Additionally, processing of AI-generated voice stimuli may differ from natural voices [32]. While this reduction was necessary for experimental control, future work is needed to understand how multi-dimensional acoustic structure influences the learning and inference in natural voices.

At present, we cannot conclusively determine whether idiosyncratic speaker representations primarily reflect long-standing differences in the perceptual representation of the voice space, differences in learning, weighting, or updating of empirical priors, or an interaction of these factors. This is difficult to resolve, as participants inevitably arrive with extensive prior auditory experience; even neonates orient toward their mother’s voice within hours of birth, suggesting prenatal tuning of voice perception [60]. Mapping individual perceptual spaces before learning speaker distributions may enable better isolation of how speaker representations are constructed relative to perceptual baseline representations.

A related consideration is whether the idiosyncratic variance estimates reflect stimulus-independent response noise, such as lapses or motor errors, rather than differences in speaker precision. Because responses are probabilistic, a less precise speaker representation can in principle produce response variability similar to that of a more precise representation with lapses. In our experiment, response consistency across stimulus repetitions cannot separate the two given the limited number of repetitions per stimulus and the absence of a ground-truth response for ambiguous test stimuli. However, participant-specific representations were stable across a one-week retest, supporting the interpretation that the idiosyncratic variances reflect persistent properties of internal representations rather than unsystematic noise. Future work could further distinguish these accounts by testing repeated trials near each speaker’s mean, where a narrow distribution predicts near-deterministic responses, so residual variability becomes attributable to noise.

## 5 Conclusion

Together, our findings show that perceptual inference in voice-identity recognition depends not only on the means of each speaker but critically also on the variance, which governs competition between multiple candidate causes under ambiguity. We demonstrate that humans learn and apply idiosyncratic, variance-sensitive speaker representations during inference that are stable over time. These findings identify precision as a key computational factor shaping perceptual inference and offer new insight into how the brain resolves uncertainty when mapping ambiguous sensory input onto internal generative models of the world.

## 6 Methods

### 6.1 Overview

Five online experiments were conducted (see Figure 3 for a schematic overview and Table 1 for detailed descriptions) to investigate whether listeners learn the mean and the variance of a speaker’s voice representation and use it as part of their internal expectations when distinguishing between competing possible speakers. In each experiment, participants learned to identify three experimentally constructed female speakers that differed in their mean, i.e., voice features (voice pitch and voice brightness), and in the variance of those features. “Voice pitch” refers here to the perceived height of the voice, which corresponds to the fundamental frequency of the sound. “Voice brightness” refers to the overall spectral quality of the voice, i.e., openness of the timbre, and was calculated by averaging the logarithms of the first three formant frequencies (F1–F3) of the vocal tract. After the learning phase, participants were presented with new, ambiguous samples and asked to identify which of the three speakers had produced the sound. Since the test samples could be closer to one or another speaker’s mean, this design allowed us to examine whether participants’ responses were consistent with Bayesian inference under competing hidden causes with different levels of associated variance. In one experiment, participants were given the option to indicate that “none of these speakers” was the cause of the utterance. This measure ensured that responses reflected a genuine perceptual inference process rather than a reliance on an explicit response strategy. From the initial learning trial to the final test trial, the average duration of the experiment was 27.47 minutes (SD = 6.44 minutes).

#### 6.1.1 Defining the speakers

To anchor the speakers in a naturalistic range, we used pooled data from four published vowel datasets [20, 29–31] to estimate the female population distribution of the voice features, specifically pitch and brightness. Because we used the pseudoword ’ada’ and chose to constrain the voice identities to sound female (mitigating expectation effects across sex), we estimated the voice space of the vowel /a/ for female speakers. Features were expressed on a logarithmic scale to account for human auditory perception [61]. This approach allowed us to create three distinct speakers that were perceptually equidistant both from the population mean and from each other in terms of pitch and brightness. We calculated a confidence ellipse (1.7 SD) around the pooled voice samples assuming no covariance. This ellipse showed that the voice identities spread much more in pitch than in voice brightness, mirroring the finding in the general population by [62], who demonstrated that a unit shift in pitch is approximately as perceptually salient as a 1.6*×* shift in brightness. In the pooled data, the pitch-to-brightness spread ratio was roughly 1.76, such that the ellipse, when represented in perceptual space, effectively forms a circle. The means of the three speakers were subsequently defined by placing points 120° apart along the perimeter of this perceptual circle. Based on a rough estimate of the natural standard deviation of voice brightness within a speaker, the standard deviations in the brightness dimension were set to 0.02, 0.03, and 0.04 log units for the low-, middle-, and high-variance conditions, respectively. To preserve the perceptual shape of the voice space, the pitch variance for each condition was calculated by multiplying the brightness variance by the perceptual scaling factor. This resulted in pitch standard deviations of approximately 0.035 (low variance), 0.053 (middle variance), and 0.07 (high variance).

Learning stimuli were then generated around the mean of the speaker according to the assigned variance level. Exemplars were arranged in concentric circles around the mean, with the number and size of circles determined by the variance condition: the low-variance condition contained one inner circle of 24 points; the middle-variance condition contained two circles (inner and middle) of 12 points each; and the high-variance condition contained three circles (inner, middle, outer) of 8 points each. Each condition therefore consisted of 24 learning exemplars per speaker.

For all analyses, we aligned speakers across participants into a common vocal reference frame with matched variance. This transformation preserved the relative configuration and variance structure of the three speaker identities while removing systematic differences in their acoustic values. Thus, standardization minimized potential confounding between the experimentally manipulated variance and speaker-specific acoustic characteristics.

#### 6.1.2 Test stimuli

Test stimuli were designed to probe participants’ inference processes across the learned voice space to assess potential perceptual effects related to differences in variance in the speaker representations. First, stimuli were sampled from a two-dimensional square grid spanning the full range of the learning exemplars in both vocal dimensions, with the grid centered on the population mean of the female voice space. To ensure that all test stimuli remained acoustically plausible, the grid was truncated by the perceptual circle used to define the voice identities, which is centered on the population mean and passes through the most extreme learning exemplars along both dimensions. This truncation prevented the generation of unnatural or implausible voice samples while preserving coverage of the relevant space.

The full square grid consisted of 11 equally spaced positions along each dimension. After truncation by the perceptual circle, the total set of test tokens included 97 points: the five central rows and columns retained all 11 points, whereas the outer three rows and columns in each direction were truncated to 9, 7, or 5 points depending on their distance from the center. This ensured a uniform sampling of the space while maintaining perceptual plausibility.

Test points were further divided into two categories based on their location within the perceptual space. Highly ambiguous test samples (21 points) fell within a circle spanning from the center of the voice space to the smallest variance circle of each speaker. These stimuli were particularly informative for assessing the influence of variance, as their categorization is expected to depend strongly on the learned variance of each speaker representation. The remaining 76 points fell outside this ambiguous region and were presented less frequently. This design allowed us to fully sample the learned voice space while emphasizing stimuli most sensitive to variance-based effects in voice identity inference.

#### 6.1.3 Example stimuli

In experiments where the variance manipulation of the speakers was explicitly communicated (Experiments 1–4), participants were presented with four example utterances for each voice identity at the start of every block to familiarize them with the to-be-learned speakers. The first example always corresponded to the mean of the respective speaker. The remaining three examples were drawn from the outermost circle of the respective voice distribution as defined by the learning stimuli, with the circle selected according to the voice-variance mapping (i.e., the inner circle for low-variance voices, the middle circle for middle-variance voices, and the outermost circle for high-variance voices). These examples were positioned equidistantly at 120° intervals along the circle, ensuring that they sampled distinct regions of the voices’ feature distribution. In the first block, these examples were located at angular positions of 7°, 127°, and 247° along the circle. In subsequent blocks, the positions were rotated in 20° increments to ensure that, across the course of the experiment, example stimuli collectively sampled the full perimeter of the respective speaker distribution.

#### 6.1.4 Voice manipulation

To ensure that voice pitch and voice brightness were the only features available for identifying a speaker, all voice samples were based on a single audio file that was manipulated. The original recording was created with the PlayHT text-to-speech synthesis platform [63] using the female voice “Marie” in German to produce the pseudoword ’ada’. The resulting .wav file was trimmed to remove silence, yielding a stimulus length of 543 ms. The acoustic analysis to situate the original sample in the voice space was conducted with Parselmouth (a Python interface to Praat [64, 65]) using custom functions. The vocal features were manipulated to match the desired values in the voice space using Praat’s “Change Gender” function via Parselmouth, shifting voice pitch to a target value and voice brightness proportionally to the ratio of target to original values. After manipulating the original voice sample, all stimuli were equalized to match an intensity of 75 dB with a Praat script [66].

### 6.2 Experimental Procedure

After providing informed consent, participants completed a headphone screening task to confirm that they could accurately perceive the auditory stimuli. If they failed on the first attempt, they were allowed one repeat. In Experiments 1–4, participants were explicitly informed that the three speakers differed in how much their vocal features varied (i.e., low, middle, or high) and were provided with four representative samples for each speaker’s voice before the start of the initial learning block and as a reminder after each test block. In Experiment 5, no explicit information about the respective variance of each speaker was provided, and thus no examples were given, so that each speaker’s variance over vocal features had to be inferred only from experience.

Across experiments, the initial learning phase contained each of the 24 stimuli from the ring-shaped learning distribution for the three speakers, resulting in 72 trials. Stimuli were presented in a pseudorandom order, ensuring that every possible transition between speakers, including repetitions, occurred equally often. On each trial, participants heard a pseudoword (“ada”) and indicated which speaker they thought had produced it by pressing one of three keys (“1”, “2”, or “3”), corresponding to numbers displayed on screen. Across experiments, participants had a maximum of 2600 ms to respond. If they failed to respond in time, a “too slow” message appeared. The mapping of each voice to a key was randomized across participants. After each response, participants received feedback. If correct, the chosen speaker number was highlighted in green; if incorrect, the correct speaker’s number remained on the screen for at least 500 ms.

After the initial learning phase, the structure of the remainder of the experiment varied across Experiments. In Experiments 1, 2, and 5, a reduced version of the initial learning and test blocks alternated separately with six learning and six test blocks. Stimulus order in test blocks was pseudorandomized so that no two consecutive trials contained stimuli adjacent in the test grid (i.e., directly above, below, or beside one another in the pitch–brightness test grid). In Experiments 3 and 4, learning and test stimuli were intermixed within a single continuous sequence, rather than being presented in separate blocks. Intermixed learning trials contained occasional feedback, while test trials were feedback-free. For these experiments, the trial sequence was fully randomized using the pandas *sample(frac=1)* function in Python [67]. Across all experiments, the same test stimuli were presented. Each of the 21 highly ambiguous stimuli was presented once per block, the mean of each speaker was presented 3 times per block, and the 76 less ambiguous stimuli were split into three groups and presented two times across the experiment. This resulted in 332 test trials in total. Similar to the initial learning, participants were asked which of the speakers most likely produced the sample and responded by pressing the respective key. In Experiment 2, participants additionally had the option to indicate that none of the speakers produced the stimulus by pressing the space key. In all test trials, a post-trial gap of 700 ms was included to allow responses that had exceeded the time limit to be recorded, although participants still saw the “too slow” message if they had failed to respond within the 2600 ms window. The reduced learning phases in Experiments 1, 2, and 5 included every second stimulus along the concentric circles around each speaker mean (e.g., 12 rather than 24 stimuli per speaker), reminding participants of the true feature distribution associated with each voice. At the end of Experiments 3, 4, and 5, participants completed a short survey rating each speaker’s voice on distinctiveness, naturalness, familiarity, masculinity, femininity, and variability using 7-point Likert scales. These parameters were later used to predict idiosyncratic variance estimates. Participants were asked to indicate the gender they believed each voice represented. Finally, they ranked the speakers according to their perceived level of variability. In Experiments 1–4, where the differences in the variance of the speakers were made explicit during learning, participants were asked both to reproduce the explicitly learned variance (88% of participants correct) and, in all experiments, to rank the three speakers from lowest to highest variance based on their perception.

### 6.3 Participants

Participants were recruited via the online platform Prolific (www.prolific.co) and were required to be native speakers of German with self-reported normal hearing. Only participants who used headphones, verified via an online headphone screening task [68], were allowed to proceed.

Each participant took part in only one of the five experiments, except in Experiment 4, where participants were intentionally recruited from Experiment 3 and invited to return approximately one week later to repeat the task. This design allowed us to assess whether any observed patterns of inference persisted across sessions or became stronger with repeated exposure.

A preregistered accuracy threshold of at least 80% correct responses (similar to [32]) per speaker during the learning blocks was required for inclusion in the analysis.

The results remained stable when including all participants. This strict criterion was preregistered to ensure that the feature distribution of each speaker was reliably acquired and could be effectively applied during the inference process for ambiguous samples. Accuracy was computed across all learning trials.

Because the acoustic features of each speaker were transformed into a common perceptual space, perceived gender was orthogonalized from the experimentally manipulated voice variance. We therefore included all participants regardless of their gender attributions to the speakers, deviating from the preregistered criterion that participants should categorize all three voices as female. In practice, approximately 52% of participants reported to have not assumed a (binary) gender for at least one of the three speakers during the task.

Including these participants preserved statistical power, while the variance-based standardization ensured that perceived gender could not systematically bias the experimental manipulation through gender-related hyperpriors.

In Experiment 1, 30 participants were recruited, of whom 18 (mean age = 28.22 years; 10 male, 8 female) met the inclusion criteria. In Experiment 2, 27 were recruited and 18 retained (mean age = 31.11 years; 6 male, 12 female). In Experiment 3, 34 were recruited and 18 retained (mean age = 31.06 years; 7 male, 11 female). In Experiment 4, 21 were recruited from Experiment 4 and 14 retained (mean age = 31.64 years; 5 male, 9 female). In Experiment 5, 169 were recruited and 108 retained (mean age = 29.95 years; 56 male, 51 female, 1 undisclosed).

### 6.4 Statistical inference

The main analyses were conducted in R using the nnet package [69] to fit multinomial logistic regression models with three response categories. The dependent variable was the participant’s speaker choice on each test trial. To interpret the model’s estimated probabilities for each response category, estimated marginal means were computed and tested using the emmeans package [70].

Our question was whether participants’ inferences about a hidden speaker identity were influenced by the variance of each speaker’s voice representation, beyond acoustic features (pitch and brightness). To test this, both responses and acoustic predictors were standardized within the voice space, such that voice category 1 corresponded to the low-variance speaker, 2 to the middle-variance speaker, and 3 to the high-variance speaker. Pitch and brightness values were mirrored and rotated into a common perceptual space across participants. This procedure ensures that any systematic preference for a particular category cannot be attributed to the specific acoustic values or the identity of the voice itself but must instead reflect an effect of speaker variance. The model can be expressed as:

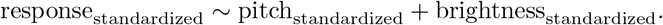

In the standardized model, the intercepts directly quantify variance-driven choice tendencies: a monotonic pattern (*β*_0,low_ *< β*_0,med_ *< β*_0,high_) indicates that, independent of acoustics, participants are progressively more likely to select the higher-variance speaker.

We deviated from the less optimal preregistered analysis plan, which involved modeling responses based on the spectral mean difference between voices and using separate variance predictors for each voice. This approach was reconsidered because (1) it does not fully eliminate voice-specific effects, and (2) the multinomial model would include variance predictors for voices irrelevant to a particular comparison (e.g., the variance of voice 3 when comparing voice 1 versus voice 2). In contrast, our standardized approach orthogonalizes variance related to the speakers, enabling a clearer and more interpretable assessment of how variance influences hidden-cause inference.

#### 6.4.1 Effect of speaker variance under stimulus uncertainty

Next, we asked whether the influence of speaker variance is amplified under greater perceptual uncertainty. We used mean reaction time (RT) as a proxy for stimulus uncertainty. This approach was preferred over the raw distance to the speaker means, since increasing distance from the center of the vocal space along the speaker direction enhances the perceived features, producing caricature-like stimuli that are easier to classify [33]. Moreover, RTs reflected a more fine-grained uncertainty structure (see Figure 4C). We validated RT as a measure of uncertainty by showing a strong correlation with mean accuracy during the learning phase (*r* = .87*, p < .*001), in which a measure of correct performance was available (in contrast to the test phase, where accuracy could not be defined since ambiguous stimuli had no objectively correct label). Accordingly, normalized reaction time was included as a continuous predictor in the standardized model:

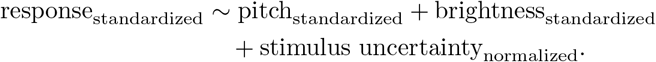

#### 6.4.2 Effect of speaker variance and experimental conditions

We examined whether the influence of speaker variances depends on the explicitness of instructions regarding speakers’ variances. Specifically, we tested for differences in effects between conditions with explicit instructions and those with implicit instructions. To this end, explicitness was incorporated as a binary categorical predictor into the multinomial model:

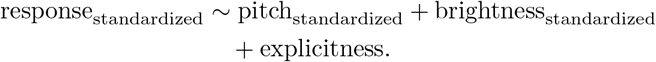

To assess whether the observed effects of speaker variances were attributable solely to response biases, we conducted an experiment in which participants had the option to respond that “none of the speakers” was the hidden cause behind an utterance. This experimental manipulation was included as an additional binary categorical predictor in the model, alongside explicitness.

If including the “none” option mitigated the effect of speaker variances, we would expect negative coefficients for this response option for the middle-and high-variance speakers, indicating a reduction in inferred causality for these speakers relative to the low-variance speaker, whose effects are primarily reflected in the intercepts. The model was specified as:

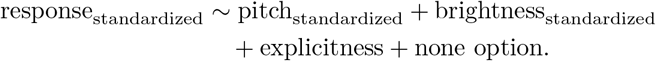

#### 6.4.3 The effect of expected variance for different learning performances

We hypothesized that participants who accurately learned the distributions of the speakers’ voices would be more likely to apply this knowledge when inferring the hidden cause of ambiguous stimuli. To test this, we examined whether the tendency to select speakers with higher voice variance as the hidden cause scaled with participants’ learning of the speakers. To avoid ceiling effects from the accuracy-based exclusion criterion, we refit the standardized model including all participants and added standardized mean learning accuracy across learning blocks as a continuous predictor:

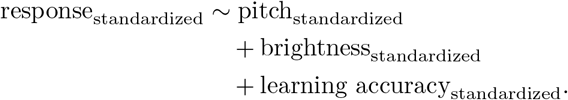

### 6.5 Computational models

#### 6.5.1 Overview

To formalize participants’ inferences about hidden speaker identity, we implemented Bayesian probabilistic speaker-attribution models in Python using PyMC (v5) [71]. The models characterize hidden-cause inference as the process by which the position of a stimulus in the two-dimensional perceptual voice space, defined by pitch and brightness, is evaluated against each speaker’s learned distribution. Speakers are represented by distributions centered on their mean, and the distance between a stimulus and each speaker’s mean determines the strength of evidence supporting that speaker as the hidden cause. These evidence values are converted into probabilistic speaker attributions via a softmax function.

We fit four variants of the model defined by systematically varying along two dimensions: whether participants learn and represent and use differences in the variance across speakers during inference, and whether variance parameters are shared across participants or estimated idiosyncratically at the participant-level. Thus, by crossing the assumptions about variance with the level of participant-specificity, we obtain four separate model variants:

1. participant-agnostic, speaker-agnostic variance;
2. participant-specific, speaker-agnostic variance;
3. participant-agnostic, speaker-specific variance; and
4. participant-specific, speaker-specific variance.

In the speaker-agnostic variance models, all speakers are assumed to share the same internal variance. This implies that participants’ inferences depend only on the distance between a stimulus and each speaker’s mean, while differences in the speaker-specific variances are disregarded. In the speaker-specific variance models, each speaker is allowed to have its own variance parameter.

This allows the computational model to capture the hypothesis that participants represent not only the mean of each voice distribution but also its variance, and that these parameters together influence the inference of the hidden cause of a stimulus.

In the participant-agnostic models, a common set of parameters is assumed across all individuals, i.e., all responses are treated as coming from a single meta-participant. In participant-specific models, idiosyncratic parameters are estimated individually for each participant using a hierarchical structure. This design allows the computational model to capture systematic differences in how participants internally represent the variance of each speaker. Here, participant-level parameters are drawn from group-level hyperpriors, allowing partial pooling that captures individual differences while still capturing group-level effects.

The fitting procedure was based on Markov chain Monte Carlo sampling algorithms. Unless stated otherwise, all models were run with four Markov chains, each with 1000 warmup samples and 1000 posterior draws, using a target acceptance probability of 0.9. Convergence was assessed via Gelman-

Rubin diagnostic *R*^ *<* 1.01 for all parameters, effective sample sizes, and visual inspection of chain traces [72]. Model comparison was performed using leave-one-out cross-validation (LOO-CV) with Pareto-smoothed importance sampling, as implemented in the ArviZ function *compare* [73].

#### 6.5.2 Model specification

The central assumption of all models is that each speaker is represented as a bivariate Gaussian distribution in a two-dimensional perceptual voice-space defined by pitch and brightness. Each stimulus is thus a point in this twodimensional space, and participants’ categorical responses are modeled as the probabilistic outcome of comparing the stimulus to the learned distributions of the three speakers.

Given a stimulus, the probability of inferring a particular speaker as the hidden cause depends on the distance of the stimulus to each speaker’s mean. For all models, the distance between a stimulus *s* and the mean *µ* of each speaker *v* is computed as the squared Euclidean distance in the two-dimensional voice space. Let the stimulus *s_i_*be:

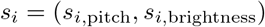

Let the mean of the distribution for each speaker be:

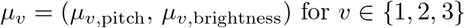

This leads to the squared Euclidean distance between stimulus *s_i_* and speaker *v* with *µ_v_*:

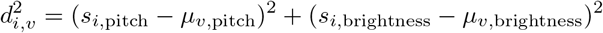

where *s_i_* is the stimulus vector and *µ_v_* is the vector of the speaker’s mean. These distances were scaled by the respective variance parameter and converted into unnormalized evidence values using a Gaussian Radial Basis Function kernel. Specifically, we defined the log evidence for a stimulus *s* given a speaker *v* as

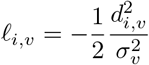

where *σ*^2^_v_ is the variance parameter for speaker *v* representing the log of the Gaussian density.

We initially considered a fully normalized Gaussian evidence function. In two dimensions, its log density includes the additional term giving:

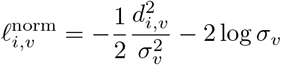

Unlike the unnormalized specification, this normalization term makes sure that all values under the curve sum up to one, thereby accounting for the effect of the scale parameter on the height of the Gaussian density. The models incorporating this normalization term with freely varying speaker-specific variance parameters exhibited sampling difficulties, including poor convergence diagnostics (*R*^^^ *>* 1.10) and low effective sample sizes (*<* 100 for 4 chains). This is likely because a difficult posterior geometry arises due to the competing influences of the distance term and the normalization term, which renders the scale parameters structurally hard to identify. We therefore used the unnormalized distance-based specification for the primary analyses because it provided a more tractable parameterization in which the scale parameters have a direct interpretation in governing the distance-dependent spread of each speaker representation.

To confirm that our theoretical conclusions hold regardless of this modeling choice, we fit a simplified normalized model in which one speaker’s scale parameter was fixed as a reference point, rendering the remaining speaker-specific scales estimable relative to this anchor. These scales were shared across participants and accompanied by a participant-specific temperature parameter to account for individual differences in response stochasticity. The model recovered the expected ordering of speaker-specific scale parameters (low *<* middle *<* high variance speaker; Supplementary Table S26) and outperformed a model with shared variance parameters, corroborating the results of the primary analyses.

Throughout the paper, we use the term precision of speaker representations to refer to the inverse of the squared scale parameter of the evidence function. Thus, smaller values for this kernel width parameter correspond to higher precision and more rapidly decreasing evidence with distance, whereas larger values correspond to broader speaker representations with more gradual decreases in evidence. The parameter was assigned a half-normal prior:

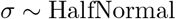

In speaker-agnostic variance models, *σ* is a single scalar shared across all speakers, either *σ*_group_ (participant-agnostic) or participant-specific with all *σ_p_* drawn from a group-level hyperprior *σ*_group_:

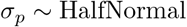

In speaker-specific variance models, *σ_v_* is allowed to differ across speakers, either shared across participants *σ_v,_*_group_ or participant-specific *σ_v,p_*with all *σ_v,p_*drawn from a group-level hyperprior *σ_v,_*_group_:

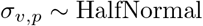

All group-level scale parameters were assigned HalfNormal priors with a scale parameter of 1.

The variance used in the likelihood computation is then given by *σ*^2^. Intuitively, a small parameter estimate for *σ* yields sharp boundaries (distance penalized strongly), whereas a large *σ* yields tolerant, broader boundaries that cover ambiguous regions. The probability of inferring speaker *v* as the hidden cause for stimulus *s_i_* is given by a softmax function over the log evidence values. The *σ* controls how flexible the softmax sharpness is per speaker. So for example in the speaker-specific variance model, for a speaker with a large *σ_v_*, even points far away from the mean still have non-negligible probability, whereas for a speaker with small *σ_v_*, the likelihood drops off quickly.

The evidence is transformed into categorical choice probabilities using a softmax function over speakers according to the formula:

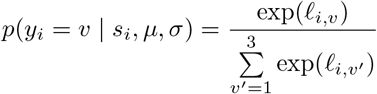

The softmax in our implementation converts the class-conditional evidence into normalized response probabilities under the assumption of uniform prior probability over the three speaker identities, rather than a maximum-likelihood decision rule. The observations were modelled via a categorical distribution using the categorical choice probabilities:

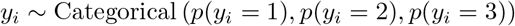

The four models were fitted on the test data of all participants (including those below the criterion) to estimate whether they used idiosyncratic variances in their speaker representations. For participants who completed the experiment a second time, only data from the first run were used for the initial fitting. The resulting winning model, i.e., the participant-specific speaker-specific variance model, was also fit to the test data from participants above the criterion only.

#### 6.5.3 Participant fits

To assess whether the speaker-specific variance structure provided a better account of behavior at the individual level, we fit the speaker-specific and speaker-agnostic variance models separately for each participant. Because these fits were performed at the individual level, no hierarchical structure was implemented. The two models were compared per participant using leave-one-out cross-validation (LOO-CV) with Pareto-smoothed importance sampling, as implemented in the ArviZ function compare [73].

#### 6.5.4 Analysis of idiosyncratic parameter estimates

To investigate how individual differences influence perception of the variance, the participant-specific, speaker-specific variance parameter estimates from the winning model were extracted and averaged over Markov chains and draws. Next, linear mixed models were fitted in R with the function *lmer* [74] on the parameter estimates as the dependent variable. To corroborate the effects of our manipulated speaker variance and the learning accuracy, the following formula was used:

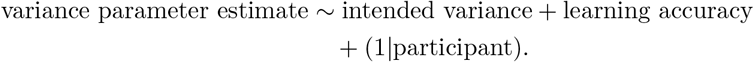

Learning accuracy was standardized, and the intended speaker variance was coded as -1, 0, and 1 for low, middle, and high.

To assess whether participant-specific speaker-specific variance estimates could be accounted for by participant characteristics or subjective perceptual differences, we extended the model to include demographic and perceptual predictors. Specifically, we examined whether participants’ age and sex, as well as subjective ratings of voice familiarity, distinctiveness, naturalness, perceived variability, maleness, and femaleness, explained variance in the estimated variance parameters. These predictors were introduced in a second modeling step for two reasons. First, this allowed us to assess whether previously established effects of intended speaker variance and learning accuracy remained robust when controlling for individual differences. Second, given the conceptual and empirical relationship between intended speaker variance and perceived variability, including subjective ratings in a subsequent step enabled us to disentangle the experimentally imposed variance from participants’ perceptual interpretations of that variance.

The extended model was specified as:

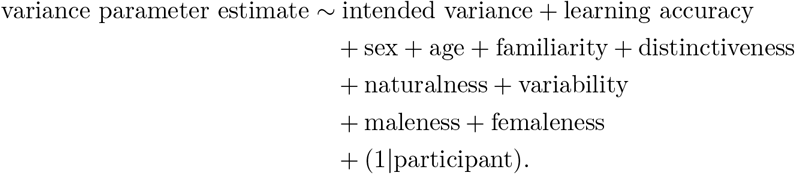

#### 6.5.5 Predicting learning accuracy

Posterior predictive simulations were conducted to assess the models’ ability to accurately reproduce each participant’s observed learning performance during the learning phase. The primary objective was to replicate the mean learning accuracy observed for each participant, effectively reversing the modeling process. We inputted the stimuli used in the learning phase into the model and generated predicted responses for each participant. Sampling was performed over chains and draws (i.e., 4,000 samples) from the posterior distributions for each participant. This process yielded predicted categorical responses. The predicted mean accuracy was then calculated as the proportion of these responses that matched the true speaker category across the posterior samples. These predicted learning accuracies were correlated with observed mean learning accuracies in the learning phase with scipy function pearsonr [75]. A high correlation demonstrates that the winning model captured individual participants’ internalized representations of the speakers.

#### 6.5.6 Cross-session prediction

A subset of participants (*n* = 21) completed the experiment a second time approximately one week later. This allowed us to test whether the internal representations of speaker-specific variances inferred by the computational model reflect stable, participant-specific inference tendencies rather than transient task-specific strategies. The underlying hypothesis was that if participants bring idiosyncratic inductive priors to the task, then model parameters estimated from a participant’s first session should predict that participant’s behavior in the second session better than parameters estimated from other participants. Conversely, if behavior were largely determined by stimulus-driven learning within the experiment, then parameters from different participants should perform similarly when predicting second-session responses. For each participant *p*, we used posterior parameter estimates obtained from fitting the winning participant-specific speaker-specific variance model to all participants first-session test data. These posterior distributions comprised participant-specific estimates of the speaker-specific variance parameters 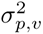. Using these parameters, we generated out-of-sample predictions for the participant’s observed choices in the second session. Specifically, for each posterior draw, we computed trial-wise categorical choice probabilities and evaluated the log-likelihood of the participant’s observed responses in the second session. Log-likelihoods were summed across all trials, yielding a distribution of total log-likelihood values per posterior draw. For each participant, responses in the second repetition were predicted using (i) the participants’ own posterior parameter estimates from the first repetition and (ii) posterior parameter estimates from every other participant. Thus, for each participant *p*, we obtained one log-likelihood distribution corresponding to *self-prediction* and N-1 distributions corresponding to *other-participant predictions*.

#### 6.5.7 Rank-based evaluation

For each prediction (self and other), log-likelihoods were averaged across posterior draws and Markov chains, yielding a single summary log-likelihood value per predicting participant. To quantify how well a participant’s own parameters predicted their second-session behavior relative to others, we ranked the self-prediction log-likelihood against the log-likelihoods obtained from all other participants’ parameters. This produced a rank score for each participant, with lower ranks indicating better predictive performance.

Under the null hypothesis that participants’ parameters are interchangeable, the expected rank of self-prediction is uniformly distributed. Deviations from this expectation indicate participant-specific stability in the inferred speaker representations. To test whether self-predictions systematically out-performed predictions based on other participants’ parameters, we compared observed self-prediction ranks to chance using a Wilcoxon signed-rank test from *scipy* [75].

#### 6.5.8 Hierarchical bayesian modeling of self–other prediction differences

To quantify the magnitude of the self-prediction advantage and account for uncertainty in the posterior estimates, we additionally modeled differences in predictive performance using a hierarchical Bayesian difference model. For each participant *p* and each other participant *q*, we computed pairwise log-likelihood differences:

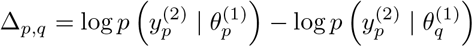

where *y*^(2)^_p_ denotes the observed responses of participant *p* in the second session, and *θ*^(1)^_p_ denotes the posterior parameter estimates obtained from the first session.

This procedure yielded a distribution of log-likelihood differences for each participant pair. These differences were aggregated across other participants, posterior draws, and chains, resulting in a participant-specific difference vector of length 21. The differences were modeled hierarchically, allowing each participant to have an individual self–other difference parameter drawn from a group-level distribution.

The group-level difference prior was specified as a normal distribution centered at zero with a scale parameter of 10, reflecting no a priori assumption of a self-prediction advantage. Group-level and participant-level scale parameters were assigned half-normal priors with scale parameters fixed at 5.

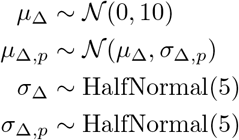

## Supplementary information

Supplementary tables are provided after the References in this manuscript.

## Declaration of Generative AI and AI-assisted Technologies

During the preparation of this work the authors used ChatGPT in order to improve language and readability. After using the tool, the authors reviewed and edited the content as needed and take full responsibility for the content.

## Acknowledgements

We thank Janika Becker, Philipp Schuhmann, Mai-Carmen Requena-Komuro, Hanna Ringer, and Henrik Eichhorn for insightful discussions, in particular regarding the figures presented in this manuscript.

## Data & Code availability

All experimental data, stimuli, and the original code used to reproduce the results and generate the figures will be publicly available on OSF upon publication.

The preregistrations connected to this manuscript can be found via the following links:

- https://osf.io/wafbh/overview?view only=5bfa5099edd441fc80df8b5d8f21f8f6
- https://osf.io/qj837/overview?view only=4141762eb6284abe8eedc0615cb58d9e

## Ethics approval

The research was approved by the local ethics committee (Ethics Committee of the Chamber of Physicians in Hamburg, approval no. PV7210).

## Author contributions

C.U.: conceptualization, methodology, software, investigation, data curation, formal analysis, visualization, writing – original draft, writing – review and editing. F.S.: methodology, software, formal analysis, writing – review and editing. H.B.: conceptualization, methodology, supervision, writing – review and editing, funding acquisition.

## Supplement

### Supplementary Tables

**Table S1.** Effect of speaker variance on hidden cause inference. Replicated results with all participants including those below-criterion. Participants’ responses (low, middle, or high variance speaker) were predicted based on the vocal features of a stimulus with a multinomial logistic regression model. This table presents the effects of stimulus vocal features (pitch and brightness) on participants’ likelihood of inferring middle or high-variance speakers, relative to a baseline (low-variance speaker). Participants were more likely to infer higher-variance speakers as a hidden cause independent of stimulus features, as reflected by significant intercepts.

| Manipulated Variance | Coefficient | Estimate | Std. Error | z-value | p-value |
| --- | --- | --- | --- | --- | --- |
| middle | (Intercept) | 0.093 | 0.012 | 7.611 | < .001 |
| middle | pitch | 4.010 | 0.028 | 142.588 | < .001 |
| middle | brightness | 0.688 | 0.025 | 27.393 | < .001 |
| high | (Intercept) | 0.265 | 0.012 | 22.798 | < .001 |
| high | pitch | 2.517 | 0.026 | 98.434 | < .001 |
| high | brightness | -2.829 | 0.026 | -110.669 | < .001 |

**Table S2.** Estimated Probabilities for hidden cause inference. Replicated results with all participants including those below-criterion. This table displays the estimated probabilities (and 95% confidence intervals) for participants’ response categories (low, middle, high variance) derived from marginal means analysis. Results indicate that participants are significantly more likely to classify utterances as coming from a high-variance speaker compared to middle or low-variance speakers, with all differences being highly significant.

| Manipulated Variance | Probability | SE | df | lower CI | upper CI | t-value | p-value |
| --- | --- | --- | --- | --- | --- | --- | --- |
| low | 0.294 | 0.002 | 6 | 0.289 | 0.299 | 135.153 | < .001 |
| middle | 0.323 | 0.002 | 6 | 0.317 | 0.328 | 145.288 | < .001 |
| high | 0.384 | 0.002 | 6 | 0.378 | 0.389 | 167.452 | < .001 |

**Table S3.** Effect of speaker variance and stimulus uncertainty on hidden cause inference. Replicated results with all participants including those below-criterion. Participants’ responses (low, middle, or high variance speaker) were predicted based on a stimulus’s vocal features and its uncertainty with a multinomial logistic regression model. This table presents the effects of vocal features (pitch and brightness) and stimulus uncertainty on participants’ likelihood of inferring middle- or high-variance speakers, relative to a baseline (low-variance speaker). Participants were more likely to infer higher-variance speakers as a hidden cause especially when stimulus uncertainty was high, as reflected by the positive and significant effect of stimulus uncertainty.

| Manipulated<br>Variance | Coefficient | Estimate | Std. Error | z-value | p-value |
| --- | --- | --- | --- | --- | --- |
| middle | (Intercept) | 0.079 | 0.013 | 5.899 | < .001 |
| middle | pitch | 4.000 | 0.028 | 140.813 | < .001 |
| middle | brightness | 0.696 | 0.025 | 27.489 | < .001 |
| middle | stim. uncertainty | 0.065 | 0.026 | 2.528 | .011 |
| high | (Intercept) | 0.250 | 0.013 | 19.793 | < .001 |
| high | pitch | 2.503 | 0.026 | 96.692 | < .001 |
| high | brightness | -2.829 | 0.026 | -110.405 | < .001 |
| high | stim. uncertainty | 0.071 | 0.025 | 2.850 | .004 |

**Table S4.** Estimated probabilities for hidden cause inference under low and high stimulus uncertainty. Replicated results with all participants including those below criterion. This table displays the estimated probabilities (and 95% confidence intervals) for participants’ response categories (low-, middle-, and high-variance speaker) under low and high stimulus uncertainty conditions (*−*1 and 1) derived from marginal means analysis. Results indicate that participants were significantly more likely to classify utterances as originating from a high-variance speaker compared to middle-or low-variance speakers when uncertainty was high, with all effects being highly significant.

| Manip.<br>Variance | Stim.<br>Uncert. | Proba-<br>bility | SE | df | Lower<br>CI | Upper<br>CI | t-ratio | p-value |
| --- | --- | --- | --- | --- | --- | --- | --- | --- |
| low | -1 | 0.311 | 0.006 | 8 | 0.297 | 0.326 | 50.547 | < .001 |
| middle | -1 | 0.316 | 0.006 | 8 | 0.301 | 0.330 | 51.365 | < .001 |
| high | -1 | 0.373 | 0.006 | 8 | 0.358 | 0.388 | 58.758 | < .001 |
| low | 1 | 0.283 | 0.004 | 8 | 0.273 | 0.292 | 68.047 | < .001 |
| middle | 1 | 0.327 | 0.004 | 8 | 0.317 | 0.337 | 73.852 | < .001 |
| high | 1 | 0.390 | 0.005 | 8 | 0.380 | 0.401 | 84.374 | < .001 |

**Table S5.** Effect of speaker variance and variance instruction on hidden cause inference. Replicated results with all participants including those below criterion. Participants’ responses (low-, middle-, or high-variance speaker) were predicted based on a stimulus’s vocal features and learning condition using a multinomial logistic regression model. This table presents the effects of vocal features (pitch and brightness) and learning instructions (explicitness; baseline = implicitness) on participants’ likelihood of inferring middle-or high-variance speakers relative to the low-variance speaker baseline. Participants were more likely to infer higher-variance speakers as the hidden cause especially when variances were instructed explicitly, as reflected by the positive and significant effect of explicitness.

| Manipulated variance | Coefficient | Estimate | Std. Error | z-value | p-value |
| --- | --- | --- | --- | --- | --- |
| middle | (Intercept) | 0.050 | 0.015 | 3.415 | .001 |
| middle | pitch | 4.012 | 0.028 | 142.574 | < .001 |
| middle | brightness | 0.686 | 0.025 | 27.286 | < .001 |
| middle | explicitness | 0.113 | 0.021 | 5.344 | < .001 |
| high | (Intercept) | 0.161 | 0.014 | 11.379 | < .001 |
| high | pitch | 2.521 | 0.026 | 98.471 | < .001 |
| high | brightness | -2.836 | 0.026 | -110.743 | < .001 |
| high | explicitness | 0.262 | 0.020 | 12.829 | < .001 |

**Table S6.** Estimated probabilities for hidden cause inference as a function of variance instruction. Replicated results with all participants including those below criterion. This table displays the estimated probabilities (and 95% confidence intervals) for participants’ response categories (low-, middle-, and high-variance speaker) under implicit and explicit variance instruction conditions derived from marginal means analysis. Results indicate that participants were significantly more likely to classify utterances as originating from a high-variance speaker compared to middle-or low-variance speakers when variance information was instructed explicitly, with all effects being highly significant.

| Manip. Variance | Instruction | Probability | SE | df | Lower CI | Upper CI | t-ratio | p-value |
| --- | --- | --- | --- | --- | --- | --- | --- | --- |
| low | implicit | 0.310 | 0.003 | 8 | 0.304 | 0.316 | 115.448 | < .001 |
| middle | implicit | 0.326 | 0.003 | 8 | 0.319 | 0.332 | 119.724 | < .001 |
| high | implicit | 0.364 | 0.003 | 8 | 0.358 | 0.371 | 130.940 | < .001 |
| low | explicit | 0.270 | 0.003 | 8 | 0.263 | 0.277 | 88.879 | < .001 |
| middle | explicit | 0.318 | 0.003 | 8 | 0.310 | 0.325 | 98.785 | < .001 |
| high | explicit | 0.412 | 0.003 | 8 | 0.405 | 0.420 | 121.191 | < .001 |

**Table S7.** Effect of speaker variance, variance instruction, and response options on hidden cause inference. Replicated results with all participants including those below criterion. Participants’ responses (low-, middle-, or high-variance speaker) were predicted based on a stimulus’s vocal features, learning instruction, and availability of a none-option using a multinomial logistic regression model. This table presents the effects of vocal features (pitch and brightness), learning instructions (explicitness; baseline = implicitness), and availability of a none-option (baseline = no none-option) on participants’ likelihood of inferring middle-or high-variance speakers relative to the low-variance speaker baseline. The effect that participants were more likely to infer higher-variance speakers as the hidden cause remained stable (or was even amplified) once the none-option was available, as indicated by the positive and significant coefficient of the none-option.

| Manipulated variance | Coefficient | Estimate | Std. Error | z-value | p-value |
| --- | --- | --- | --- | --- | --- |
| middle | (Intercept) | 0.050 | 0.015 | 3.415 | .001 |
| middle | pitch | 4.013 | 0.028 | 142.577 | < .001 |
| middle | brightness | 0.686 | 0.025 | 27.288 | < .001 |
| middle | explicit | 0.078 | 0.023 | 3.368 | .001 |
| middle | none-option | 0.150 | 0.039 | 3.854 | < .001 |
| high | (Intercept) | 0.161 | 0.014 | 11.381 | < .001 |
| high | pitch | 2.522 | 0.026 | 98.475 | < .001 |
| high | brightness | -2.836 | 0.026 | -110.740 | < .001 |
| high | explicit | 0.233 | 0.022 | 10.480 | < .001 |
| high | none-option | 0.124 | 0.037 | 3.325 | .001 |

**Table S8.** Estimated probabilities for hidden cause inference as a function of none-option availability. Replicated results with all participants including those below criterion. This table displays the estimated probabilities (and 95% confidence intervals) for participants’ response categories (low-, middle-, and high-variance speaker) when the none-option was available or not, derived from marginal means analysis. Results indicate that participants were significantly more likely to classify utterances as originating from a high-variance speaker compared to middle-or low-variance speakers when the none-option was available, with all effects being highly significant.

| Manip. Variance | None Option | Proba-bility | SE | df | Lower CI | Upper CI | t-ratio | p-value |
| --- | --- | --- | --- | --- | --- | --- | --- | --- |
| low | no | 0.293 | 0.002 | 10 | 0.288 | 0.298 | 125.456 | < .001 |
| middle | no | 0.320 | 0.002 | 10 | 0.314 | 0.325 | 133.205 | < .001 |
| high | no | 0.387 | 0.003 | 10 | 0.382 | 0.393 | 154.756 | < .001 |
| low | yes | 0.266 | 0.006 | 10 | 0.252 | 0.279 | 42.898 | < .001 |
| middle | yes | 0.337 | 0.007 | 10 | 0.322 | 0.352 | 49.633 | < .001 |
| high | yes | 0.397 | 0.007 | 10 | 0.382 | 0.413 | 57.235 | < .001 |

**Table S9.** Effect of speaker variance on hidden cause inference (above-criterion sample). Participants’ responses (low-, middle-, or high-variance speaker) were predicted based on the vocal features of a stimulus using a multinomial logistic regression model. This table presents the effects of stimulus vocal features (pitch and brightness) on participants’ likelihood of inferring middle-or high-variance speakers relative to the low-variance speaker baseline. Participants were more likely to infer higher-variance speakers as the hidden cause independent of stimulus features, as reflected by significant intercepts.

| Manipulated variance | Coefficient | Estimate | Std. Error | z-value | p-value |
| --- | --- | --- | --- | --- | --- |
| middle | (Intercept) | 0.23 | 0.02 | 12.67 | < .001 |
| middle | pitch | 4.94 | 0.04 | 117.22 | < .001 |
| middle | brightness | 0.67 | 0.04 | 18.51 | < .001 |
| high | (Intercept) | 0.38 | 0.02 | 22.05 | < .001 |
| high | pitch | 3.33 | 0.04 | 86.11 | < .001 |
| high | brightness | -3.76 | 0.04 | -96.54 | < .001 |

**Table S10.**
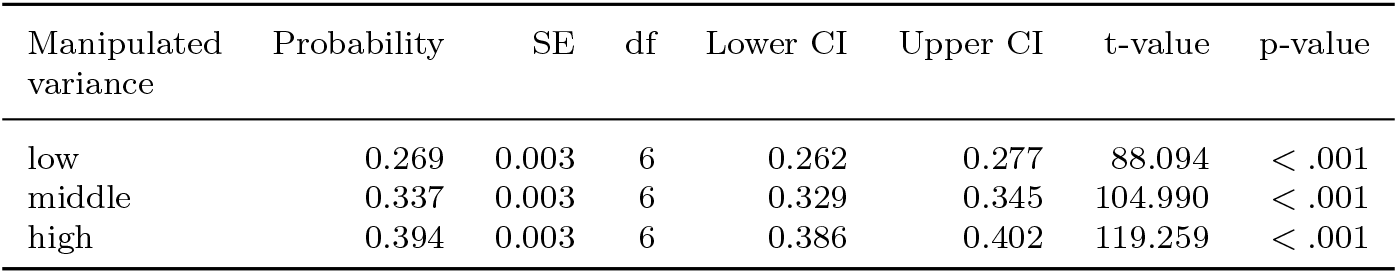
Estimated probabilities for hidden cause inference (above-criterion sample). This table displays the estimated probabilities (and 95% confidence intervals) for participants’ response categories (low-, middle-, and high-variance speaker) derived from marginal means analysis. Results indicate that participants were significantly more likely to classify utterances as originating from a high-variance speaker compared to middle-or low-variance speakers, with all differences being highly significant.

| Manipulated variance | Probability | SE | df | Lower CI | Upper CI | t-value | p-value |
| --- | --- | --- | --- | --- | --- | --- | --- |
| low | 0.269 | 0.003 | 6 | 0.262 | 0.277 | 88.094 | < .001 |
| middle | 0.337 | 0.003 | 6 | 0.329 | 0.345 | 104.990 | < .001 |
| high | 0.394 | 0.003 | 6 | 0.386 | 0.402 | 119.259 | < .001 |

**Table S11.** Effect of speaker variance and stimulus uncertainty on hidden cause inference (above-criterion sample). Participants’ responses (low-, middle-, or high-variance speaker) were predicted based on a stimulus’s vocal features and uncertainty using a multinomial logistic regression model. This table presents the effects of vocal features (pitch and brightness) and stimulus uncertainty on participants’ likelihood of inferring middle-or high-variance speakers relative to the low-variance speaker baseline. Participants were more likely to infer higher-variance speakers as the hidden cause especially when stimulus uncertainty was high, as reflected by the positive and significant effect of stimulus uncertainty.

| Manipulated variance | Coefficient | Estimate | Std. Error | z-value | p-value |
| --- | --- | --- | --- | --- | --- |
| middle | (Intercept) | 0.191 | 0.020 | 9.404 | < .001 |
| middle | pitch | 4.901 | 0.043 | 115.101 | < .001 |
| middle | brightness | 0.693 | 0.037 | 18.787 | < .001 |
| middle | stim. uncertainty | 0.121 | 0.037 | 3.246 | < .001 |
| high | (Intercept) | 0.328 | 0.020 | 16.680 | < .001 |
| high | pitch | 3.294 | 0.039 | 83.797 | < .001 |
| high | brightness | -3.764 | 0.039 | -96.231 | < .001 |
| high | stim. uncertainty | 0.180 | 0.037 | 4.899 | < .001 |

**Table S12.** Estimated probabilities for hidden cause inference under low and high stimulus uncertainty (above-criterion sample). This table displays the estimated probabilities (and 95% confidence intervals) for participants’ response categories (low-, middle-, and high-variance speaker) under low and high stimulus uncertainty conditions (*−*1 and 1) derived from marginal means analysis. Results indicate that participants were significantly more likely to classify utterances as originating from a high-variance speaker compared to middle-or low-variance speakers when uncertainty was high, with all effects being highly significant.

| Manip.<br>Variance | Stim.<br>Uncert. | Proba-<br>bility | SE | df | Lower<br>CI | Upper<br>CI | t-ratio | p-value |
| --- | --- | --- | --- | --- | --- | --- | --- | --- |
| low | −1 | 0.309 | 0.009 | 8 | 0.287 | 0.331 | 32.835 | < .001 |
| middle | −1 | 0.332 | 0.009 | 8 | 0.310 | 0.353 | 35.534 | < .001 |
| high | −1 | 0.359 | 0.010 | 8 | 0.338 | 0.381 | 37.806 | < .001 |
| low | 1 | 0.248 | 0.005 | 8 | 0.236 | 0.260 | 46.833 | < .001 |
| middle | 1 | 0.339 | 0.006 | 8 | 0.325 | 0.353 | 56.027 | < .001 |
| high | 1 | 0.413 | 0.006 | 8 | 0.398 | 0.428 | 64.530 | < .001 |

**Table S13.** Effect of speaker variance and variance instruction on hidden cause inference (above-criterion sample). Participants’ responses (low-, middle-, or high-variance speaker) were predicted based on a stimulus’s vocal features and learning condition using a multinomial logistic regression model. This table presents the effects of vocal features (pitch and brightness) and learning instructions (explicitness; baseline = implicitness) on participants’ likelihood of inferring middle-or high-variance speakers relative to the low-variance speaker baseline. Participants were more likely to infer higher-variance speakers as the hidden cause especially when variances were instructed explicitly, as reflected by the positive and significant effect of explicitness.

| Manipulated<br>variance | Coefficient | Estimate | Std. Error | z-value | p-value |
| --- | --- | --- | --- | --- | --- |
| middle | (Intercept) | 0.189 | 0.021 | 9.128 | < .001 |
| middle | pitch | 4.940 | 0.042 | 117.207 | < .001 |
| middle | brightness | 0.673 | 0.036 | 18.448 | < .001 |
| middle | explicit | 0.097 | 0.029 | 3.338 | .001 |
| high | (Intercept) | 0.293 | 0.020 | 14.438 | < .001 |
| high | pitch | 3.333 | 0.039 | 86.128 | < .001 |
| high | brightness | -3.768 | 0.039 | -96.572 | < .001 |
| high | explicit | 0.220 | 0.029 | 7.664 | < .001 |

**Table S14.** Estimated probabilities for hidden cause inference as a function of variance instruction (above-criterion sample). This table displays the estimated probabilities (and 95% confidence intervals) for participants’ response categories (low-, middle-, and high-variance speaker) under implicit and explicit variance instruction conditions derived from marginal means analysis. Results indicate that participants were significantly more likely to classify utterances as originating from a high-variance speaker compared to middle-or low-variance speakers when variance instruction was explicit, with all effects being highly significant.

| Manip. Variance | Variance Instr. | Probability | SE | df | Lower CI | Upper CI | t-ratio | p-value |
| --- | --- | --- | --- | --- | --- | --- | --- | --- |
| low | implicit | 0.281 | 0.004 | 8 | 0.273 | 0.290 | 76.732 | < .001 |
| middle | implicit | 0.340 | 0.004 | 8 | 0.331 | 0.349 | 88.352 | < .001 |
| high | implicit | 0.378 | 0.004 | 8 | 0.369 | 0.387 | 95.904 | < .001 |
| low | explicit | 0.250 | 0.004 | 8 | 0.240 | 0.259 | 59.594 | < .001 |
| middle | explicit | 0.333 | 0.005 | 8 | 0.322 | 0.343 | 72.015 | < .001 |
| high | explicit | 0.418 | 0.005 | 8 | 0.407 | 0.429 | 85.757 | < .001 |

**Table S15.** Effect of speaker variance, variance instruction, and response options on hidden cause inference (above-criterion sample). Participants’ responses (low-, middle-, or high-variance speaker) were predicted based on a stimulus’s vocal features, learning instruction, and availability of a none-option using a multinomial logistic regression model. This table presents the effects of vocal features (pitch and brightness), learning instructions (explicitness; baseline = implicitness), and availability of a none-option (baseline = no none-option) on participants’ likelihood of inferring middle-or high-variance speakers relative to the low-variance speaker baseline. The effect that participants were more likely to infer higher-variance speakers as the hidden cause remained stable (or slightly increased) once the none-option was available, as indicated by the small positive coefficient of the none-option (significant only for the high-variance category).

| Manipulated variance | Coefficient | Estimate | Std. Error | z-value | p-value |
| --- | --- | --- | --- | --- | --- |
| middle | (Intercept) | 0.189 | 0.021 | 9.126 | < .001 |
| middle | pitch | 4.940 | 0.042 | 117.203 | < .001 |
| middle | brightness | 0.673 | 0.036 | 18.448 | < .001 |
| middle | explicitness | 0.103 | 0.032 | 3.201 | .001 |
| middle | none option | -0.021 | 0.053 | -0.404 | .686 |
| high | (Intercept) | 0.293 | 0.020 | 14.429 | < .001 |
| high | pitch | 3.333 | 0.039 | 86.120 | < .001 |
| high | brightness | -3.769 | 0.039 | -96.577 | < .001 |
| high | explicitness | 0.192 | 0.032 | 6.075 | < .001 |
| high | none option | 0.105 | 0.051 | 2.051 | .040 |

**Table S16.** Estimated probabilities for hidden cause inference as a function of none-option availability (above-criterion sample). This table displays the estimated probabilities (and 95% confidence intervals) for participants’ response categories (low-, middle-, and high-variance speaker) when the none-option was available or not, derived from marginal means analysis. Results indicate that participants were significantly more likely to classify utterances as originating from a high-variance speaker compared to middle-or low-variance speakers when the none-option was available, with all effects being highly significant.

| Manip.<br>Variance | None<br>Option | Proba-<br>bility | SE | df | Lower<br>CI | Upper<br>CI | t-ratio | p-value |
| --- | --- | --- | --- | --- | --- | --- | --- | --- |
| low | no | 0.267 | 0.003 | 10 | 0.259 | 0.274 | 81.194 | < .001 |
| middle | no | 0.339 | 0.004 | 10 | 0.331 | 0.347 | 96.379 | < .001 |
| high | no | 0.394 | 0.004 | 10 | 0.386 | 0.402 | 108.578 | < .001 |
| low | yes | 0.257 | 0.008 | 10 | 0.239 | 0.276 | 31.260 | < .001 |
| middle | yes | 0.320 | 0.009 | 10 | 0.300 | 0.340 | 36.007 | < .001 |
| high | yes | 0.423 | 0.010 | 10 | 0.401 | 0.444 | 44.069 | < .001 |

**Table S17.**
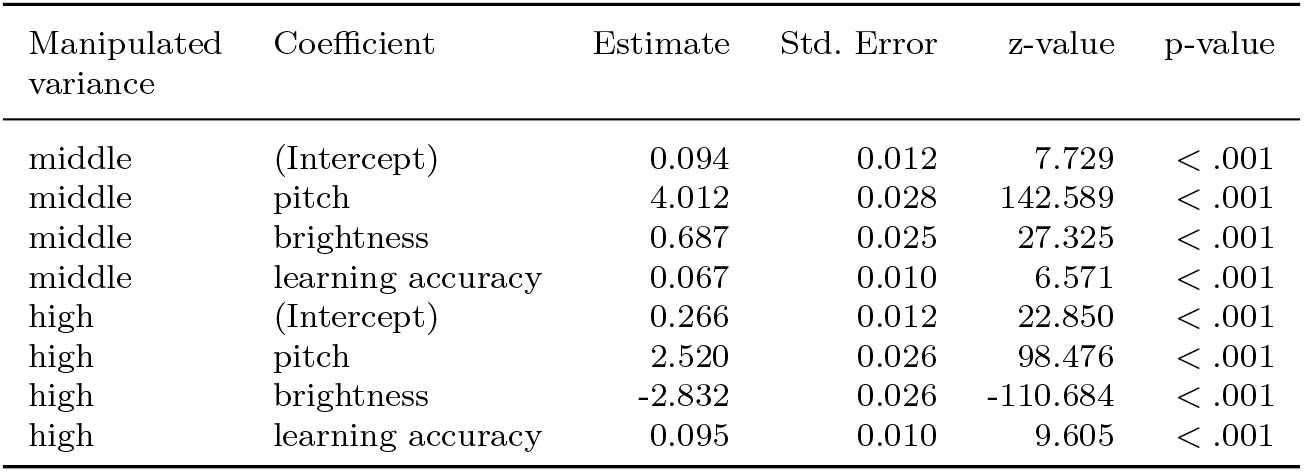
Effect of speaker variance and learning accuracy on hidden cause inference. Participants’ responses (low-, middle-, or high-variance speaker) were predicted based on a stimulus’s vocal features and participants’ learning accuracy using a multinomial logistic regression model. This table presents the effects of vocal features (pitch and brightness) and learning accuracy on participants’ likelihood of inferring middle-or high-variance speakers relative to the low-variance speaker baseline. Participants were more likely to infer higher-variance speakers as the hidden cause when learning accuracy was higher, as reflected by the positive and significant coefficients of learning accuracy.

| Manipulated<br>variance | Coefficient | Estimate | Std. Error | z-value | p-value |
| --- | --- | --- | --- | --- | --- |
| middle | (Intercept) | 0.094 | 0.012 | 7.729 | < .001 |
| middle | pitch | 4.012 | 0.028 | 142.589 | < .001 |
| middle | brightness | 0.687 | 0.025 | 27.325 | < .001 |
| middle | learning accuracy | 0.067 | 0.010 | 6.571 | < .001 |
| high | (Intercept) | 0.266 | 0.012 | 22.850 | < .001 |
| high | pitch | 2.520 | 0.026 | 98.476 | < .001 |
| high | brightness | -2.832 | 0.026 | -110.684 | < .001 |
| high | learning accuracy | 0.095 | 0.010 | 9.605 | < .001 |

**Table S18.** Estimated probabilities for hidden cause inference as a function of learning accuracy. This table displays the estimated probabilities (and 95% confidence intervals) for participants’ response categories (low-, middle-, and high-variance speaker) when learning accuracy was 1 SD below (77.8%) and 1 SD above (95.3%) the mean learning accuracy, derived from marginal means analysis. Results indicate that participants were significantly more likely to classify utterances as originating from a high-variance speaker compared to middle-or low-variance speakers when learning accuracy was higher, with all effects being highly significant.

| Manip. Variance | Learning Accuracy | Prob. | SE | df | Lower CI | Upper CI | t-ratio | p-value |
| --- | --- | --- | --- | --- | --- | --- | --- | --- |
| low | 0.953 | 0.277 | 0.003 | 8 | 0.270 | 0.283 | 99.596 | < .001 |
| middle | 0.953 | 0.325 | 0.003 | 8 | 0.319 | 0.332 | 110.423 | < .001 |
| high | 0.953 | 0.398 | 0.003 | 8 | 0.391 | 0.405 | 129.224 | < .001 |
| low | 0.778 | 0.311 | 0.003 | 8 | 0.305 | 0.318 | 107.147 | < .001 |
| middle | 0.778 | 0.320 | 0.003 | 8 | 0.313 | 0.326 | 108.487 | < .001 |
| high | 0.778 | 0.369 | 0.003 | 8 | 0.362 | 0.376 | 121.047 | < .001 |

**Table S19.** Model comparison for above-criterion participants. Models are ordered from best (rank 0) to worst (rank 3). The participant-specific speaker-specific variance model is identified as the best-fitting model. ELPD-LOO: expected log pointwise predictive density estimated via leave-one-out cross-validation; higher values indicate better out-of-sample predictive performance. p-LOO: effective number of parameters. ELPD diff: difference in ELPD-LOO relative to the best model. Weight: model weight. SE: standard error of the ELPD-LOO estimate. SE diff: standard error of the ELPD-LOO difference relative to the best model.

| Model | Rank | ELPD-LOO | p-LOO | ELPD diff | Weight | SE | SE diff |
| --- | --- | --- | --- | --- | --- | --- | --- |
| speaker-specific participant-specific | 0 | -32574.143 | 579.552 | 0.000 | 0.847 | 179.584 | 0.000 |
| speaker-agnostic participant-specific | 1 | -33747.522 | 187.338 | 1173.379 | 0.000 | 179.502 | 60.035 |
| speaker-specific participant-agnostic | 2 | -34003.207 | 3.791 | 1429.064 | 0.048 | 181.621 | 71.109 |
| speaker-agnostic participant-agnostic | 3 | -34217.042 | 1.253 | 1642.899 | 0.104 | 182.346 | 74.642 |

**Table S20.** Group-level estimates for the standard deviation (*σ*) parameter of each speaker for above-criterion participants. The variance is the square of the standard deviation (*σ*^2^). The parameter ordering corresponds to the intended speaker variance (low, middle, high). Mean: posterior mean estimate of *σ*. SD: posterior standard deviation. HDI 3% and 97%: lower and upper bounds of the highest density interval. MCSE mean and MCSE SE: Monte Carlo standard error of the mean and standard deviation estimates, respectively. ESS (bulk and tail): effective sample size for the bulk and tails of the posterior distribution, indicating sampling efficiency.

| Manipulated variance | Mean | SD | HDI 3% | HDI 97% | MCSE mean | MCSE SE | ESS bulk | ESS tail | $\hat{R}$ |
| --- | --- | --- | --- | --- | --- | --- | --- | --- | --- |
| $\sigma$ low | 0.447 | 0.025 | 0.401 | 0.496 | < .001 | < .001 | 5547 | 3035 | 1.00 |
| $\sigma$ middle | 0.467 | 0.026 | 0.422 | 0.518 | < .001 | < .001 | 5479 | 3027 | 1.00 |
| $\sigma$ high | 0.498 | 0.028 | 0.447 | 0.552 | < .001 | < .001 | 5720 | 3422 | 1.00 |

**Table S21.** Model comparison for the full dataset. Models are ordered from best (rank 0) to worst (rank 3). The participant-specific speaker-specific variance model is identified as the best-fitting model. ELPD-LOO: expected log pointwise predictive density estimated via leave-one-out cross-validation; higher values indicate better out-of-sample predictive performance. p-LOO: effective number of parameters. ELPD diff: difference in ELPD-LOO relative to the best model. Weight: model weight. SE: standard error of the ELPD-LOO estimate. SE diff: standard error of the ELPD-LOO difference relative to the best model.

| Model | Rank | ELPD-LOO | p-LOO | ELPD<br>diff | Weight | SE | SE<br>diff |
| --- | --- | --- | --- | --- | --- | --- | --- |
| speaker-specific<br>participant-specific | 0 | −59575.463 | 895.353 | 0.000 | 0.886 | 215.288 | 0.000 |
| speaker-agnostic<br>participant-specific | 1 | −61370.843 | 295.321 | 1795.380 | 0.037 | 215.132 | 73.242 |
| speaker-specific<br>participant-agnostic | 2 | −63943.094 | 3.448 | 4367.631 | 0.000 | 217.731 | 109.961 |
| speaker-agnostic<br>participant-agnostic | 3 | −64162.740 | 1.184 | 4587.277 | 0.078 | 218.265 | 112.276 |

**Table S22.**
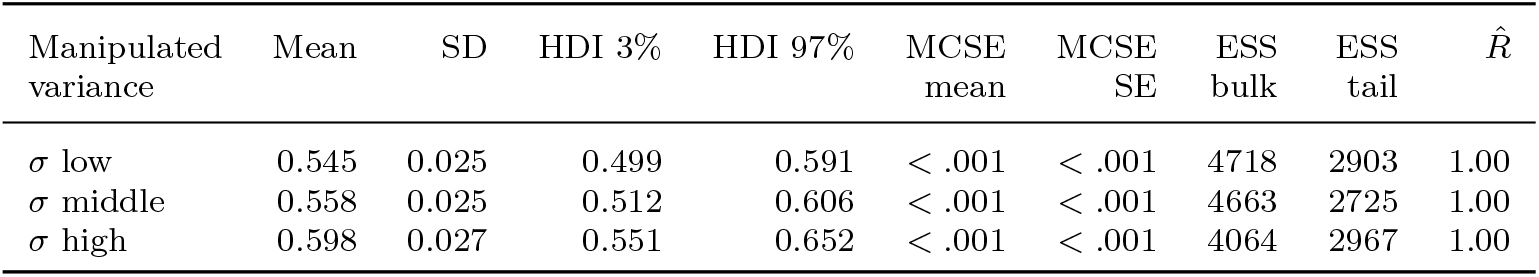
Group-level estimates for the standard deviation *σ* parameter of each speaker (full dataset). The variance is the square of the standard deviation (*σ*^2^). The parameter ordering corresponds to the intended speaker variance (low, middle, high). Mean: posterior mean estimate of *σ*. SD: posterior standard deviation. HDI 3% and 97%: lower and upper bounds of the highest density interval. MCSE mean and MCSE SE: Monte Carlo standard error of the mean and standard deviation estimates. ESS (bulk and tail): effective sample size for bulk and tail posterior samples, indicating sampling efficiency.

| Manipulated<br>variance | Mean | SD | HDI 3% | HDI 97% | MCSE<br>mean | MCSE<br>SE | ESS<br>bulk | ESS<br>tail | $\hat{R}$ |
| --- | --- | --- | --- | --- | --- | --- | --- | --- | --- |
| $\sigma$ low | 0.545 | 0.025 | 0.499 | 0.591 | < .001 | < .001 | 4718 | 2903 | 1.00 |
| $\sigma$ middle | 0.558 | 0.025 | 0.512 | 0.606 | < .001 | < .001 | 4663 | 2725 | 1.00 |
| $\sigma$ high | 0.598 | 0.027 | 0.551 | 0.652 | < .001 | < .001 | 4064 | 2967 | 1.00 |

**Table S23.** Fixed effect coefficients from the linear mixed-effects model predicting idiosyncratic speaker-specific variance estimates. The model indicates that intended speaker variance (coded as low = *−*1, middle = 0, high = 1) is significantly positively associated with variance estimates, whereas learning accuracy shows a significant negative association.

| Coefficient | Estimate | Std. Error | df | t-value | p-value |
| --- | --- | --- | --- | --- | --- |
| (Intercept) | 0.534 | 0.005 | 258 | 98.171 | < .001 |
| Intended variance | 0.027 | 0.004 | 519 | 5.976 | < .001 |
| Learning accuracy | −0.131 | 0.005 | 258 | −24.051 | < .001 |

**Table S24.** Results of the linear mixed-effects model including subjective perception predictors of idiosyncratic speaker-specific variance estimates. The analysis shows that, beyond intended speaker variance and learning accuracy, perceived voice variability significantly positively predicts variance estimates. Listener age shows a significant negative association, whereas other subjective perception measures do not reach statistical significance (*p > .*05).

| Coefficient | Estimate | Std. Error | df | t-value | p-value |
| --- | --- | --- | --- | --- | --- |
| (Intercept) | 0.434 | 0.036 | 190.908 | 12.008 | < .001 |
| Intended variance | 0.012 | 0.005 | 388.416 | 2.396 | .017 |
| Learning accuracy | -0.124 | 0.006 | 194.406 | -20.091 | < .001 |
| Listener sex | 0.013 | 0.012 | 190.520 | 1.080 | .281 |
| Listener age | 0.003 | 0.001 | 191.278 | 2.442 | .015 |
| Perceived variability | 0.013 | 0.005 | 522.590 | 2.873 | .004 |
| Perceived naturalness | -0.008 | 0.006 | 547.992 | -1.400 | .162 |
| Perceived distinctiveness | -0.007 | 0.004 | 517.070 | -1.628 | .104 |
| Perceived familiarity | -0.008 | 0.006 | 554.164 | -1.484 | .138 |
| Perceived maleness | 0.001 | 0.006 | 565.897 | 0.183 | .855 |
| Perceived femaleness | 0.002 | 0.006 | 552.406 | 0.268 | .789 |

**Table S25.** Results of the hierarchical Bayesian model estimating the self-prediction advantage. Positive parameter estimates indicate a self-prediction advantage compared to other participants. The table reports the group-level parameter and individual participant estimates.

| Parameter | Mean | SD | HDI 3% | HDI 97% | MCSE<br>mean | MCSE<br>SE | ESS<br>bulk | ESS<br>tail | $\hat{R}$ |
| --- | --- | --- | --- | --- | --- | --- | --- | --- | --- |
| Group difference | 30.73 | 2.74 | 26.03 | 36.12 | 0.03 | 0.02 | 9044 | 2837 | 1.00 |
| Group SD | 12.77 | 1.68 | 9.72 | 15.92 | 0.02 | 0.01 | 9789 | 3093 | 1.00 |
| Participant SD | 26.23 | 2.05 | 22.42 | 29.98 | 0.02 | 0.01 | 11246 | 3017 | 1.00 |
| **Individual difference** |  |  |  |  |  |  |  |  |  |
| Participant 1 | 23.52 | 0.04 | 23.44 | 23.60 | 0.00 | 0.00 | 11023 | 2218 | 1.00 |
| Participant 2 | 15.11 | 0.03 | 15.06 | 15.16 | 0.00 | 0.00 | 10576 | 3167 | 1.00 |
| Participant 3 | 26.01 | 0.04 | 25.94 | 26.08 | 0.00 | 0.00 | 12496 | 2687 | 1.00 |
| Participant 4 | 48.80 | 0.04 | 48.72 | 48.87 | 0.00 | 0.00 | 11966 | 2792 | 1.00 |
| Participant 5 | 15.89 | 0.03 | 15.83 | 15.96 | 0.00 | 0.00 | 11426 | 2698 | 1.00 |
| Participant 6 | 21.26 | 0.03 | 21.20 | 21.32 | 0.00 | 0.00 | 9965 | 2994 | 1.00 |
| Participant 7 | 49.64 | 0.04 | 49.57 | 49.70 | 0.00 | 0.00 | 13363 | 2529 | 1.00 |
| Participant 8 | 29.45 | 0.03 | 29.39 | 29.51 | 0.00 | 0.00 | 10555 | 2540 | 1.00 |
| Participant 9 | 25.48 | 0.05 | 25.39 | 25.57 | 0.00 | 0.00 | 9993 | 2703 | 1.00 |
| Participant 10 | 22.36 | 0.03 | 22.30 | 22.42 | 0.00 | 0.00 | 13141 | 2799 | 1.00 |
| Participant 11 | 38.82 | 0.04 | 38.73 | 38.90 | 0.00 | 0.00 | 11385 | 3065 | 1.00 |
| Participant 12 | 47.40 | 0.03 | 47.34 | 47.44 | 0.00 | 0.00 | 12620 | 2948 | 1.00 |
| Participant 13 | 30.78 | 0.03 | 30.72 | 30.85 | 0.00 | 0.00 | 11512 | 2555 | 1.00 |
| Participant 14 | 23.53 | 0.04 | 23.45 | 23.61 | 0.00 | 0.00 | 10272 | 2465 | 1.00 |
| Participant 15 | 43.55 | 0.04 | 43.48 | 43.63 | 0.00 | 0.00 | 11459 | 3115 | 1.00 |
| Participant 16 | 33.36 | 0.04 | 33.28 | 33.44 | 0.00 | 0.00 | 12497 | 2677 | 1.00 |
| Participant 17 | 55.19 | 0.04 | 55.11 | 55.27 | 0.00 | 0.00 | 10257 | 2993 | 1.00 |
| Participant 18 | 52.25 | 0.04 | 52.17 | 52.32 | 0.00 | 0.00 | 8841 | 2770 | 1.00 |
| Participant 19 | 10.00 | 0.03 | 9.94 | 10.06 | 0.00 | 0.00 | 10339 | 2479 | 1.00 |
| Participant 20 | 29.73 | 0.06 | 29.63 | 29.84 | 0.00 | 0.00 | 11599 | 2570 | 1.00 |
| Participant 21 | 53.98 | 0.04 | 53.91 | 54.06 | 0.00 | 0.00 | 9929 | 2498 | 1.00 |

**Table S26.** Group-level estimates for the standard deviation (*σ*) parameter of each speaker with normalized log evidence (above-criterion participants). Results from a simplified, normalized model in which the scale parameter for the low-variance speaker was fixed at 0.1 as a reference point, and the remaining speaker-specific scale parameters were estimated relative to it. Scale parameters were shared across participants, while a participant-specific temperature parameter captured individual differences in response stochasticity. The model recovered the expected ordering of speaker-specific scale parameters (low-variance *<* middle-variance *<* high-variance speaker). Mean: the mean estimated value for *σ*. SD: the standard deviation of the estimated *σ*. HDI 3% and 97%: lower and upper bound of the highest density interval. MCSE mean and MCSE SE: Markov Chain Monte Carlo Standard Error estimates for the mean and the standard error of the mean; they indicate the precision of the MCMC estimates. ESS (Effective Sample Size): estimated number of independent samples drawn from the posterior distribution, for the bulk (ESS bulk) and the tails (ESS tail), indicating sampling efficiency.

| Speaker | Mean | SD | HDI 3% | HDI 97% | MCSE<br>mean | MCSE<br>SE | ESS<br>bulk | ESS<br>tail | $\hat{R}$ |
| --- | --- | --- | --- | --- | --- | --- | --- | --- | --- |
| $\sigma$ low | 0.100 | 0.000 | 0.100 | 0.100 | < .001 | < .001 | 4000 | 4000 | — |
| $\sigma$ middle | 0.112 | 0.001 | 0.111 | 0.113 | < .001 | < .001 | 3790 | 3178 | 1.00 |
| $\sigma$ high | 0.119 | 0.001 | 0.118 | 0.120 | < .001 | < .001 | 3936 | 3395 | 1.00 |

